# Distinct tumor immune microenvironmental (TIME) landscapes drive divergent immunotherapy responses in glioblastoma

**DOI:** 10.1101/2025.10.18.683203

**Authors:** Linqian Weng, Krish Skandha Gopalan, Mélanie Guyot, Maxime Vanmechelen, Pouya Nazari, Marie Duhamel, Clelia Donisi, Carla Pallarés-Moratalla, Abhishek D. Garg, Kieron White, Annette T. Byrne, Diether Lambrechts, Frederik De Smet, Gabriele Bergers

## Abstract

**Background:** Immunotherapies have improved outcomes in many cancers but show limited efficacy in glioblastoma (GBM). This study aimed to determine whether immunotherapy could be tailored to GBM by functional subtyping of vascular-immune landscapes.

**Methods:** We employed single-cell RNA sequencing, multiplex immunohistochemistry to characterize three distinct TIME subtypes in human and murine GBMs. We evaluated responses to combination of anti-angiogenic immunomodulating therapies (CD40 Ag, anti-PDL1, PI3Kγ/δ inhibition) in orthotopic syngeneic GBM mouse models.

**Results:** We identified three distinct functional TIME subtypes with unique vascular-immune landscapes in human and murine GBM: TIME-low (immune-low/deserted, leaky vasculature), TIME-med (intermediate immune-infiltration, angiogenic), and TIME-high (heavily infiltrated with immunosuppressive myeloid cells and dysfunctional T cells). Representative mouse models of TIME-GBMs responded in subtype specific ways to anti-angiogenic immunomodulating therapies. TIME-low GBMs enhanced T-cell activity but relapsed due to emerging myeloid immunosuppression, concomitant with mesenchymal transition. TIME-med displayed the most immune-activated, yet angiogenic phenotype, and showed overall good responses to various anti-angiogenic immunomodulating therapies. TIME-high GBMs were mostly non-responsive but improved when the myeloid-cell PI3Kγ was targeted. However, CD40 agonist treatment, expected to enhance APC function, unexpectedly worsened survival by promoting angiogenesis and heightening immunosuppression, leading to dysfunctional T cells and reduced NK cell recruitment, and subsequent enhanced tumor propagation.

**Conclusions:** Our study reveals three GBM TIME subtypes with distinct vascular-immune landscapes that require tailored therapies. TIME-med tumors are predicted to respond best to immunotherapies, TIME-low tumors show transient effects with anti-angiogenic immunomodulating therapies, while TIME-high tumors, due to their profound immunosuppression, can even have worse outcomes.

**Key points:**

- Three TIME subtypes were identified in GBM with distinct vascular-immune landscapes
- TIME subtypes show divergent immunotherapy responses
- TIME classification supports personalized treatment strategy for GBM immunotherapy

**Importance of the Study:** This study advances glioblastoma immunotherapy by providing the first comprehensive single-cell characterization of TIME subtypes, moving beyond bulk RNA-sequencing to reveal detailed functional states of immune cells. We establish clinically relevant murine models that recapitulate human TIME subtypes, enabling preclinical testing of TIME-targeted therapies. Our findings identify TIME-low GBM as immune deserted and TIME-med tumors as the most immunotherapy-responsive subtype that should be prioritized for clinical selection. We found that high immune infiltration correlates with non-responsiveness and even unexpected detrimental effects with CD40 agonist treatment in TIME-high tumors—critical information given ongoing clinical trials. Identifying distinct immunosuppressive mechanisms across TIME subtypes and differential treatment responses provides a framework for personalized immunotherapy selection. The immediate translational impact of this work highlights the importance of TIME classification for treatment stratification and the urgent need to consider TIME status in clinical trial design, potentially explaining variable patient responses in previous GBM immunotherapy trials.

## Introduction

Glioblastoma (GBM), the most aggressive adult brain tumor, exhibits relentless growth and resistance to therapy^1^. Despite intensive efforts, the standard of care has remained largely unchanged for nearly two decades, highlighting the urgent need for innovative therapeutic strategies.

With the increasing recognition of the tumor immune microenvironment (TIME) in regulating tumor progression, targeting both the vasculature (via anti-angiogenic therapies) and immune cells (via immune checkpoint blockade, ICB) have shown promise in solid cancers^2–7^. However, while these strategies showed early potential, their translation to GBM has yielded limited clinical success. Recurrent GBMs are generally unresponsive to ICB-based therapies, with only a few patients showing survival benefits^8^. Bevacizumab, an FDA-approved anti-VEGFA antibody, has improved radiographic responses and quality of life but has shown minimal benefit for overall survival, benefitting only a subset of patients in the neoadjuvant setting^9–12^. Similarly, combining ICB with anti-angiogenic therapy has not significantly outperformed monotherapy, with benefits restricted to certain patient subgroups^13,14^. This selective responsiveness has driven interest in better characterizing the immune heterogeneity of GBM. To date, most efforts have relied on bulk RNA-seq data and cellular deconvolution methods to infer immune cell abundance. While informative, these approaches lack single-cell resolution and are often confounded by noise and complex statistical assumptions. Only recently have GBMs been classified into tumor microenvironment (TME)-based subtypes: TME-low (immune desert), TME-med (immune-heterogeneous with angiogenic features), and TME-high (high leukocyte infiltration with some immunotherapeutic responsiveness)^15^. Despite these advances, bulk RNA-seq remains largely limited in its ability to resolve individual cell states^16^. The emergence of scRNA-seq has enabled a more precise, cell-level view of the TIME^17^. In GBM, scRNA-seq has revealed new immune programs, intercellular signaling, and potential therapeutic targets^18–21^. However, these studies have largely focused on single immune lineages rather than comprehensive TIME classification.

In this study, we recapitulate the vascular-immune landscapes of GBM at single-cell resolution across the three established TIME subtypes. We identify distinct functional immune states: TIME-low tumors are immune-deserted, TIME-med tumors are both immunogenic and angiogenic, and TIME-high tumors exhibit strong immunosuppressive features. We further identify syngeneic, orthotopic preclinical models that reflect these TIME subtypes and evaluate their response to immunomodulatory therapies. These include anti-angiogenic immunotherapies and CD40 agonists, which demonstrate variable—and in some cases adverse—responses depending on TIME subtype. Our findings reveal the intricate interplay of immune and vascular elements in GBM and emphasize the need for tailored TIME subtype-specific treatment strategies.

## Materials and Methods

### Tumor models and animal studies

NFpp10-GFP GBM cells were derived from C57BL/6 neural stem cells (NSCs) with knockdown of *Trp53*, *Nf1*, and *Pten*. NSCG-mCherry cells originated from Ink4/Arf−/− NSCs with ectopic EGFRvIII expression. For intracranial transplantation, 6-8 week-old FvB/N or C57BL/6 mice were anesthetized and mounted in a stereotactic apparatus. Tumor cells (5K NSCG-mCherry or 20K NFpp10-GFP) were injected into the right striatum and grown for 14 days before analysis.

Treatment regimens included:

- **BP** (5 mg/kg B20 + 10 mg/kg anti-PD-L1)
- **BPI** (BP + 10 mg/kg IPI-145)
- **CD40/DC vaccine** (100 μg CD40 agonist/dose)

Drugs were administered intraperitoneally every 3 days for BP/BPI and every 5 days for CD40 agonist, starting 6–7 days post-implantation. Survival experiments continued until neurological symptoms or humane endpoints. Brains were collected for analysis.

### Immunofluorescence

Tumors were fixed in 2% PFA, cryoprotected in 30% sucrose, and sectioned at 10 μm. Sections were stained with antibodies against various markers. For vessel functionality, TMR-dextran and tomato lectin were injected intravenously before sacrifice. Images were acquired using Zeiss microscopes (20×) and analyzed using ImageJ and QuPath. CD31-based pixel classifiers aided vessel segmentation.

### Flow cytometry

Single-cell suspensions were prepared using mechanical and enzymatic dissociation (Tumor Dissociation Kit, gentleMACS Dissociator). Cells were stained with antibody panels targeting T cells, myeloid cells, NK cells, and functional markers. Viability dye excluded dead cells. Data were collected on BD Fortessa and analyzed in FlowJo.

### MILAN multiplex immunohistochemistry

Tissue microarrays were constructed from 2–6 representative 2-mm tumor cores. The MILAN protocol involved sequential rounds of 3-marker + DAPI staining, high-definition imaging, and antibody stripping (SDS/β-mercaptoethanol). This cycle repeated across multiple marker combinations.

Bioinformatic analysis using DISSCOvery included preprocessing, image registration, background removal, and DAPI-based segmentation. Cell identities were inferred using six core markers (CD1C, CD3, CD4, CD8, SOX2, CD68, TMEM119) with k-means and Phenograph clustering. Additional markers (FOXP3, LAG3, PDCD1, KI67, CD163, HLA-DR) defined functional states.

### TIME subgrouping

For MILAN and scRNA-seq datasets, cell proportions were normalized and clustered using partition around medoids (Manhattan distance, k=3). Elbow method and bootstrap validation (n=1000) confirmed cluster stability (Jaccard > 0.65).

### Single-cell RNA sequencing and analysis

Tumors were digested using collagenases and DNase, and single-cell suspensions processed with 10X Genomics Chromium (targeting 6,000 cells/library). Libraries were sequenced on Illumina HiSeq4000 and aligned to the mouse genome (mm10) using CellRanger v7.0.1.

Data were processed in Seurat4 with lenient filtering to retain low-transcription cells (e.g., neutrophils). Doublets were removed (scDblFinder), data normalized, and batch effects corrected using Harmony. PCA and UMAP were used for dimensionality reduction, with clustering via FindNeighbors and FindClusters.

SCENIC (Nextflow) inferred transcription factor networks. Enrichment analysis used GO, Reactome, and Hallmark pathways, scored by AUCell. Differential gene expression used Wilcoxon Rank Sum test (Seurat’s FindMarkers). Visualizations were generated using custom R packages (e.g., SeuratExtend^22^).

### Statistical analysis

Statistical tests (t-test, Log-rank, Mann-Whitney, Wilcoxon) were performed using Prism and R. Significance was defined as *p*<0.05 with multiple comparison corrections as needed. Further details are provided in the Supplemental Material.

## Results

### Human GBM displays distinct tumor-immune microenvironment subtypes

To reveal TIME subtypes at single-cell resolution, we encompassed a cohort of 52 primary IDH-wildtype GBM samples from 35 patients (**Fig. 1A**). Although several scRNA-seq studies of glioma exist, with recent efforts shifting towards the generation of large-scale integrative pan-glioma atlases^23^, we observed that per-patient representation of non-malignant cells (by proportion of sequenced cells) was irregular across datasets. To uniformly characterize the TME on a patient level, we opted for a conservative approach, curating 6 droplet-based (10X Genomics) scRNA-seq datasets and used a consistent preprocessing methodology with a unified reference genome, followed by downstream integrative analysis. After quality control, we identified 240,625 high-quality cells, comprising five host cell types (Myeloid, Lymphocytes, Oligodendrocytes, Endothelial Cells (EC), and Pericytes), besides tumor cells (**Fig. 1B**, **S1A**). Validating results previously shown in bulk RNA-seq analyses^15,24,25^, we observed significant variation in immune cell abundance across samples (**Fig. S1B**), confirming intertumoral heterogeneity.

**Figure 1.**
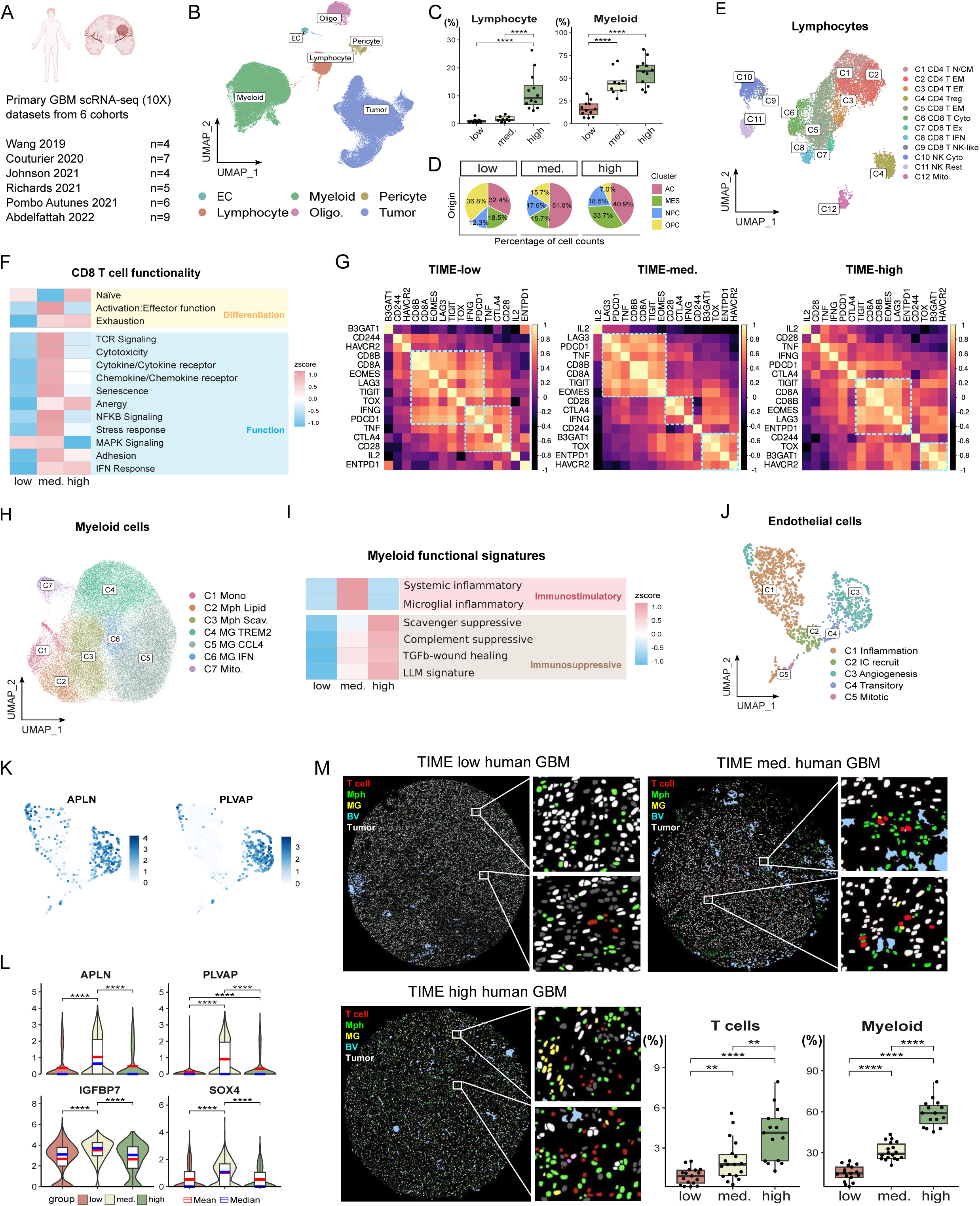
Characterization of immune microenvironmental subtypes in human GBM. (A) Schematic representation of primary GBM scRNA-seq (10X) datasets from 6 cohorts used in this study. (B) UMAP plot showing major cell types identified in human GBM samples. (C) Quantification of lymphocyte and myeloid cell proportions across TIME-low, TIME-intermediate (med.), and TIME-high GBM samples. (D) Distribution of GBM molecular subtypes across TIME-low, TIME-med, and TIME-high samples. (E) UMAP plot of lymphocyte subpopulations in human GBM. (F) Heatmap showing differential expression of CD8+ T cell functional signatures across TIME-low, TIME-med., and TIME-high samples, grouped by differentiation and function categories. (G) Correlation analysis of “chronically sensitized” CD8+ T cell signature in TIME-low, TIME-med, and TIME-high samples. (H) UMAP plot of myeloid cell subpopulations in human GBM. (I) Heatmap showing expression of myeloid functional signatures across TIME subtypes, categorized as immunostimulatory and immunosuppressive. (J) UMAP plot of endothelial cell subpopulations in human GBM. (K) Feature plots showing expression of APLN and PLVAP in endothelial cells. (L) Violin plots showing expression of APLN, PLVAP, IGFBP7, and SOX4 across TIME subtypes in endothelial cells. (M) Representative MILAN multiplex immunohistochemistry images of TIME-low, TIME-med, and TIME-high human GBM samples, and quantification T cell and myeloid cell proportions across different subtypes. The images show T cells (red), macrophages (Mph, yellow), microglia (MG, green), blood vessels (BV, blue), and tumor cells (white). Statistical significance: *p < 0.05, **p < 0.01, ***p < 0.001, ****p < 0.0001.

We applied partition-around-medoid (PAM) clustering on the relative proportions of immune cells, delineating three subgroups with varying TIME composition, as described before^15^: TIME-low, corresponding to tumors with little immune infiltration, TIME-high, corresponding to tumors significantly enriched in both myeloid and lymphoid cells, and TIME-med, with immune infiltration intermediate between these two extremes (**Fig. 1C**). The TIME subtypes contained all malignant gene programs^26^ at varying percentages with TIME-low GBM more skewed towards an oligodendrocyte-progenitor-like (OPC-like) program, TIME-med tumors enriched in the astrocyte-like (AC-like) program, and TIME-high tumors with a higher prevalence of the mesenchymal-like (MES-like) program (**Fig. 1D**).

### Human GBM subtypes reveal distinct immune functional status

We next characterized the functional status of the lymphoid compartment in the TIME subtypes. Unbiased subcluster analysis revealed a diverse landscape of lymphoid populations including several subtypes of CD4^+^ T cells, CD8^+^ T cells, and NK cells (**Fig. 1E**). Examining differences in subcluster proportion across TIME subtypes, TIME-high tumors had a greater abundance of regulatory T cells (Tregs), while TIME-med tumors were more enriched CD8^+^ T cells/NK cells (**Fig. S1C**). Utilizing a set of curated, pan-cancer T cell gene signatures^27^, enrichment analysis of differentiation and function-related pathways revealed subtype-specific patterns in both CD4^+^ and CD8^+^ T cell compartments. T cells in TIME-low tumors exhibited a relative lack of differentiation and functional pathway enrichment, consistent with a lymphocyte-deserted phenotype, but T cells in TIME-med tumors were concordantly enriched in nearly all functional pathways, displaying elevated TCR signaling, cytotoxicity, and cytokine/chemokine pathways. In contrast, T cells in TIME-high tumors upregulated the expression of gene signatures implicated in exhaustion, anergy, and Tregs, indicating functional impairment (**Fig. 1F**, **S1D**). Cross-correlation analysis of a pan-cancer “chronically-sensitized” CD8^+^T-cell (csCD8+T) signature as a readout of CD8^+^T-cell activity^28^, using diffusion mapping^29^ revealed that TIME-low and especially TIME-med tumors had a relatively greater concordance within the signature-associated genes, partially indicative of a more “traditional” tumor sensitization phenotype, while TIME-high tumors had a low concordance, indicative of a severely dysfunctional phenotype (**Fig. 1G**). Notably, increased concordance has been linked to immunotherapy response, suggesting that TIME-med (and to some extent TIME-low) tumors could show better clinical outcomes when treated with ICB, an observation that challenges prior indications that higher immune infiltration is associated with an ICB-response phenotype^15^, but might be in line with observations of some T cell ‘pseudo-hot’ tumors being ICB-resistant^30^.

Analysis of the myeloid compartment in the three TIME subtypes identified seven distinct myeloid subpopulations: monocytes (Mono), mitotic cells (Mito.), lipid-associated macrophages (Lipid Mph), scavenger macrophages (Scav Mph), and TREM2-expressing microglia (TREM2 MG), as well as CCL4-expressing microglia (CCL4 MG), interferon-inflamed microglia (IFN MG) (**Fig. 1H**). Notably, we independently found subclusters (i.e., CCL4 MG and Scav Mph) that mirrored activity programs (‘microglial inflammatory’ and ‘scavenger immunosuppressive’, respectively) in a pan-glioma analysis of myeloid cells^31^. Of note, TIME-med tumors harbored the highest proportion of CCL4 MG cells, while displaying reduced frequencies of Lipid Mph and Scav Mph cells (**Fig. S1E**). TIME-med myeloid cells expressed elevated levels of pro-inflammatory genes (e.g., *TNF*, *CX3CR1*, *CCL3*, *CCL4*), whereas TIME-high myeloid cells expressed higher levels of anti-inflammatory genes (e.g., *SPP1*, *MRC1*, *FN1*, *VEGFA*, and *CD163*) (**Fig. S1F**). In line with the concept that myeloid cells may have varied functional immune status, we compiled both immune-stimulating and immunosuppressive gene expression programs of myeloid cells in GBM^28,31,32^. Enrichment analysis of the myeloid signatures revealed a lack of enrichment in TIME-low tumors, congruent with the low abundance of myeloid cells, TIME-med GBMs were highly enriched for immunostimulatory signatures, and myeloid cells in TIME-high GBMs were enriched for multiple immunosuppressive signatures (**Fig.1I**). A key observation here is that though TIME-high tumors in general are more immunosuppressive, there is no single putative myeloid cell state that associated with this immunosuppression. Myeloid-derived immunosuppression encompasses multiple functional pathways, as evidenced by enrichment of not only glioma-intrinsic immunosuppressive pathways (i.e., ‘Scavenger suppressive’ and ‘Complement suppressive’), but also exhibits enrichment of generic immunosuppression mediated by lipid-laden and TGFβ-wound healing-like pathways, a testament to their phenotypic plasticity.

Moreover, we turned our attention to the vasculature, given that immune infiltration into tumors is regulated by tumor ECs and that GBMs are considered highly angiogenic. Using markers from a prior GBM EC study^33^, we identified five transcriptionally distinct EC subclusters (**Fig. 1J**). Interestingly, angiogenic ECs exhibited high expression of *APLN* (apelin) and *PLVAP*, known to promote vessel permeability and leakiness^33^(**Fig. 1K**), reminiscent of an angiogenic and leaky vasculature in GBM^33,34^. TIME-low tumors contained the highest proportion of inflammatory endothelial cells, potentially influencing immune cell trafficking. TIME-med tumors exhibited a significantly higher proportion of the angiogenic ECs compared to both TIME-low and TIME-high subtypes, suggesting a unique vascular activation state in this group (**Fig. S1G**). In addition to *APLN* and *PLVAP*, ECs in TIME-med tumors showed elevated expression of *IGFBP7* and *SOX4* (**Fig. 1L**), genes known to be associated with angiogenesis and metabolic reprogramming^35–37^. This expression profile suggests that TIME-med tumors may sustain a more metabolically active and angiogenic vasculature.

### Validation of TIME subtypes in human GBM via spatial proteomics

As a validation cohort, we aimed to recapitulate our TIME subtyping on the protein level, utilizing the MILAN (Multiple Iterative Labeling by Antibody Neodeposition) high-resolution multiplex immunohistochemistry platform, in 50 primary, IDH-WT human GBM samples. We again applied PAM-based clustering to the relative proportions of immune cells, identifying TIME-low, TIME-med, and TIME-high subtypes that are analogous to those identified via scRNA-seq (**Fig. 1M**). Examination of the vasculature suggested that TIME-med tumors exhibited a trend towards increased vessel density, aligning with our earlier findings of enhanced angiogenic activity (**Fig. S1H**). While TIME-high tumors displayed greater infiltration of CD4^+^ and CD8^+^ T cells, they also had an increased abundance of Tregs, resulting in a slight elevation of the Treg/CD8 T cell ratio (**Fig. S1I**). In contrast, both TIME-low and TIME-med tumors tended to have more proliferating T cells, as well as higher frequencies of PD1^+^ and LAG3^+^ T cells compared to TIME-high tumors (**Fig. S1J**), indicating a greater pool of potentially ICB re-activable and tumor-sensitized lymphocytes. Myeloid cell analysis similarly revealed a trend toward increased CD163^+^ macrophages in TIME-low and TIME-med tumors, though not statistically significant due to heterogeneity within TIME-high samples (**Fig. S1K**). Notably, TIME-med tumors also displayed a trend towards more MHCII+ myeloid cells and proliferating myeloid subsets, suggesting enhanced antigen presentation capacity (**Fig. S1K-L**). Correlation analysis of immune cell abundances across subtype, revealed significantly stronger intra-tumoral concordance in TIME-med tumors compared to TIME-low and TIME-high groups (**Fig. S1M**), indicative of more coordinated immune responses in the TIME-med context.

Collectively, these data reinforce the distinct immune profiles across the three TIME subtypes. While TIME-low are considered immune-deserted or low, TIME-med GBM emerge as a more functionally active subtype, with proliferative, checkpoint-expressing T cells and immunostimulatory myeloid components. In contrast, TIME-high tumors, albeit marked by high infiltration, exhibit an immune dysfunctional and anergic phenotype with versatile suppressive myeloid cells.

### Murine tumor models recapitulate human GBM subtypes

The differing functional immune states of the TIME subtypes offer prognostic insight and may inform immunotherapeutic strategies for GBM. Thus, we set out to identify representative preclinical murine models that recapitulated their phenotypic variation. Based on prior literature confirming immunogenicity through extensive immunophenotyping^38,39^, we recognized the GL261 model—one of the most widely used syngeneic models of GBM—as a potential candidate for the TIME-med subtype. However, this should be interpreted with caution, as the tumor is carcinogen-induced and therefore hypermutated, a phenomenon rarely observed in human GBM patients^40^. Further, we noted that the immune index of the genetically engineered NSCG and NFpp10 GBMs was representative of a TIME-low and TIME-high GBM, respectively. NSCG tumors^41,42^ originate from *Arf/Ink4a*-deficient neural stem cells harboring EGFRvIII, while NFpp10 tumors^4,43,44^ derive from neural stem cells are deficient in *Nf1*, *p53*, and *Pten* (**Fig. 2A**).

**Figure 2.**
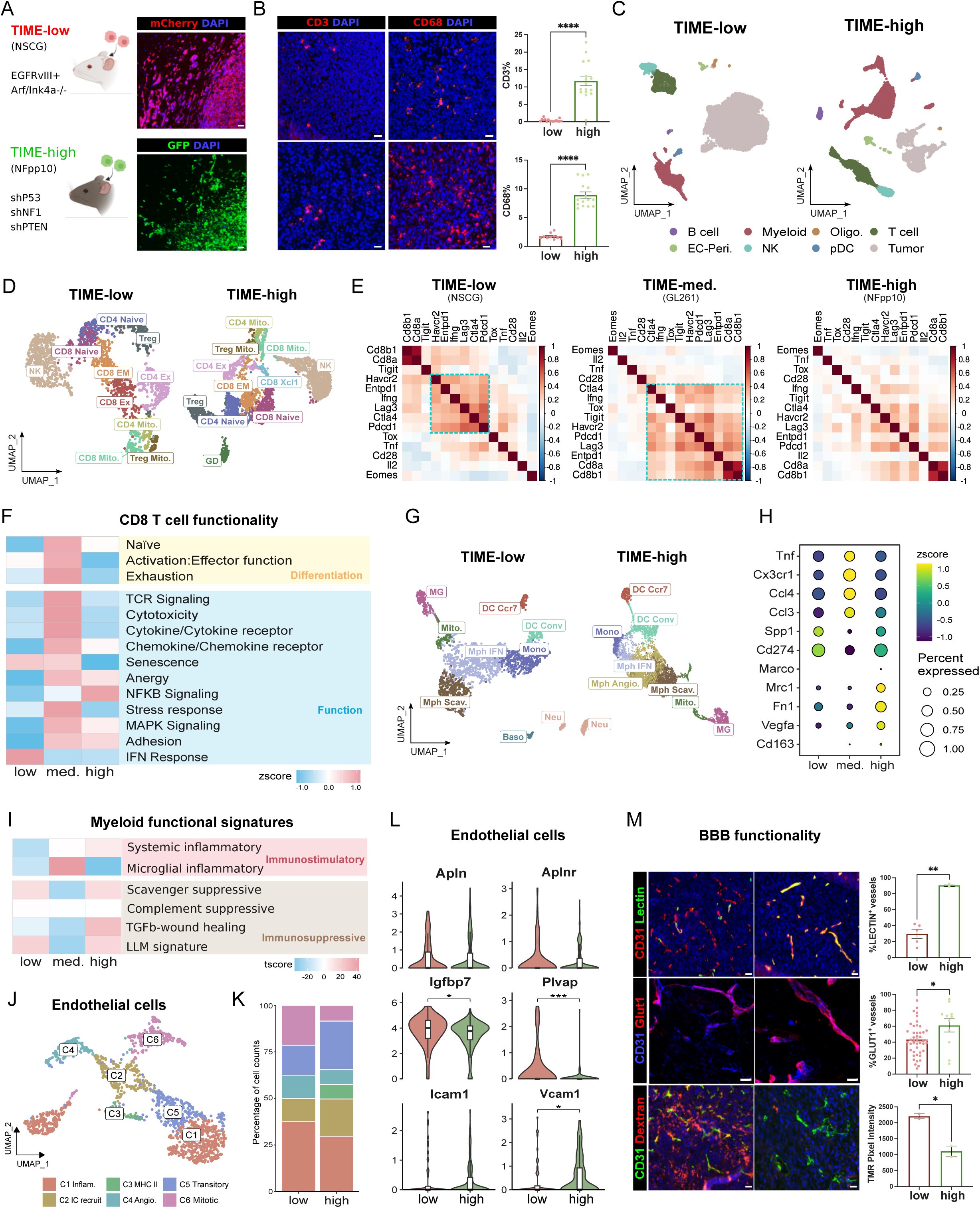
Murine GBM models recapitulate human GBM subtypes. (A) Schematic representation of TIME-low (NSCG) and TIME-high (NFpp10) murine GBM models with their key genetic alterations. (B) Representative immunofluorescence (IF) images and quantification of CD3^+^ and CD68^+^ immune cells in TIME-low and TIME-high tumors. (C) UMAP plots showing major cell types identified in TIME-low and TIME-high murine GBM models by scRNA-seq. (D) UMAP plots of T and NK cell subpopulations in TIME-low and TIME-high tumors. (E) Correlation analysis of “chronically sensitized” CD8+ T cell signature in TIME-low, TIME-med, and TIME-high murine tumors. (F) Heatmap showing differential expression of CD8+ T cell functional signatures across TIME-low, TIME-med, and TIME-high murine tumors, grouped by differentiation and function categories. (G) UMAP plots of myeloid cell subpopulations in TIME-low and TIME-high tumors. (H) Dot plot showing expression of inflammatory and immunosuppressive markers across TIME subtypes. Color indicates expression level (z-score), and dot size represents percentage of cells expressing the gene. (I) Heatmap showing expression of myeloid functional signatures across TIME subtypes, categorized as immunostimulatory and immunosuppressive. (J) UMAP plot of murine GBMs endothelial cell subpopulations. (K) Relative composition (%) of endothelial cell subclusters across TIME-low and TIME-high tumors. (L) Violin plots showing expression of endothelial cell markers in TIME-low and TIME-high tumors. (M) Representative IF images and quantification of vascular perfusion (lectin) and blood-brain barrier (BBB) integrity markers (Glut1, dextran) in TIME-low and TIME-high tumors. Data are presented as mean ± SEM. Statistical significance: *p < 0.05, **p < 0.01, ***p < 0.001, ****p < 0.0001. (Mann-Whitney test or as indicated).

Immunofluorescence and flow cytometry confirmed substantial immune infiltration in NFpp10 (CD45^+^ cells, CD3^+^ T cells, macrophages, microglia, and higher CD4^+^/CD8^+^ T-cell counts), contrasting with the sparse immune presence in NSCG (**Fig. 2B**, **S2A-B**). We therefore considered NSCG as TIME-low, and NFpp10 as TIME-high GBM, while GL261 exhibited intermediate infiltration patterns^45–47^, supporting its TIME-med classification. ScRNA-seq of 8,147 cells from NSCG (TIME-low) and 7,500 cells from NFpp10 (TIME-high) tumors revealed conserved cell populations across both models, recapitulating the cellular landscape observed in human GBM (**Fig. 2C**). Subtype analysis inferred a higher prevalence of MES-like identity in TIME-high tumors, whereas TIME-low tumors exhibited mixed MES, AC, and NPC-like profiles (**Fig. S2C**).

We next assessed the immune cell functionality in the different murine TIME subtypes. Unsupervised clustering revealed distinct T-cell and NK subpopulations (**Fig. 2D**). Integration of publicly available GL261 CD45^+^ scRNA-seq data confirmed that TIME-med CD8+ T cells expressed both high cytotoxic effector genes (*Prf1*, *Gzmb*, *Nkg7*, *Ifng*) and immune inhibitory receptors (*Pdcd1*, *Ctla4*, *Tigit*, *Lag3*), paralleling the functional signature of human TIME-med tumors (**Fig. S2D**). Correlation analyses mirrored human trends: TIME-med tumors had the strongest concordance for the genes within the csCD8^+^T cell-signature, whereas TIME-high tumors showed the weakest, suggesting more severe CD8^+^ T-cell dysfunction in the latter (**Fig. 2E**). Functional pathway analysis further indicated more active, immune-responsive T cells in TIME-med tumors, implying greater immunotherapy responsiveness (**Fig. 2F**, **S2E**), suggesting greater potential to respond to immunotherapy.

Myeloid compartment profiling identified conserved populations across TIME-low and TIME-high tumors, including monocytes, microglia, macrophage subsets (IFN-related, scavenger, angiogenic), dendritic cells (conventional, Ccr7+), and neutrophils (**Fig. 2G**). Scavenger macrophages, enriched in *Arg1*, *Mrc1* (CD206), and lipid-laden macrophage (LLM) signature (**Fig. S2F**), indicated strong immunosuppressive activity. TIME-med TAMs expressed inflammatory markers (*Tnf*, *Cx3cr1*, *Ccl3*, *Ccl4*) and myeloid inflammatory pathways, whereas TIME-high TAMs upregulated immunosuppressive markers (*Marco*, *Mrc1*, *Fn1*, *Vegfa*, *Cd163*) and TGFβ–wound-healing pathways (**Fig. 2H-I**). Cross-species correlation confirmed strong alignment between murine and human myeloid subclusters (**Fig. S2G**).

EC analysis revealed murine GBM EC populations analogous to human counterparts. We also identified an EC subset uniquely enriched in TIME-high tumors that expressed high levels of MHC class II molecules (**Fig. 2J**). While *Apln* and *Aplnr* expression was similar across TIME-low and TIME-high, the former exhibited elevated *Igfbp7* and *Plvap*, and harbored more angiogenic and mitotic ECs, consistent with aberrant angiogenesis (**Fig. 2K-L**). TIME-high ECs expressed lower *Igfbp7* and *Plvap* but higher adhesion molecules (*Icam1* and *Vcam1*) (**Fig. 2L**), suggesting superior lymphocyte recruitment potential. Functionally, TIME-low vasculature displayed poor perfusion (lectin), reduced BBB integrity (Glut1), and increased permeability (dextran) compared to the vasculature of TIME-high GBM, indicative of a more severe BBB disruption, possibly contributing to low immune infiltration (**Fig. 2M**).

Collectively, our comprehensive analysis of lymphocytes, myeloid cells, and vascular cells in different TIME subtypes recapitulates the distinct immune and vascular features of human TIME subtypes. This validation underscores their translational relevance and highlights immune vulnerabilities that may be exploited for subtype-specific therapeutic development.

### TIME-low, but not TIME-high GBM, respond to anti-angiogenic ICB

Given the distinct vascular and immune states in TIME subtypes, we examined how anti-angiogenic immunotherapy responses correlate with TIME status. TIME-med tumors, with their immunologically active and angiogenic profiles, are predicted to respond well, as supported by studies showing GL261/TIME-med GBMs benefit from anti-angiogenic, ICB, and combination therapies^40,48,49^. We tested anti-angiogenic ICB therapy in TIME-low/NSCG and TIME-high/NFpp10 GBM using a-VEGF/B20 (murine bevacizumab) plus aPD-L1 (BP; **Fig. 3A**), as PD-L1 is widely expressed in myeloid, endothelial, and tumor cells (**Fig. S3A-B**) and is upregulated after anti-VEGF therapy^3,4^. In TIME-low GBM, BP significantly improved survival, yielding ~25% long-term survivors, increased tumor apoptosis, and greater CD3^+^ T cell and macrophage infiltration (**Fig. 3B-C**). In contrast, TIME-high tumors showed no survival benefit, minimal apoptosis, and reduced CD3^+^ infiltration. Flow cytometry confirmed TIME-low tumors exhibited increased CD4^+^, CD8^+^, and Gzmb^+^ CD8^+^ T cells and a higher CD8/Treg ratio, unlike TIME-high tumors (**Fig. 3D**).

**Figure 3.**
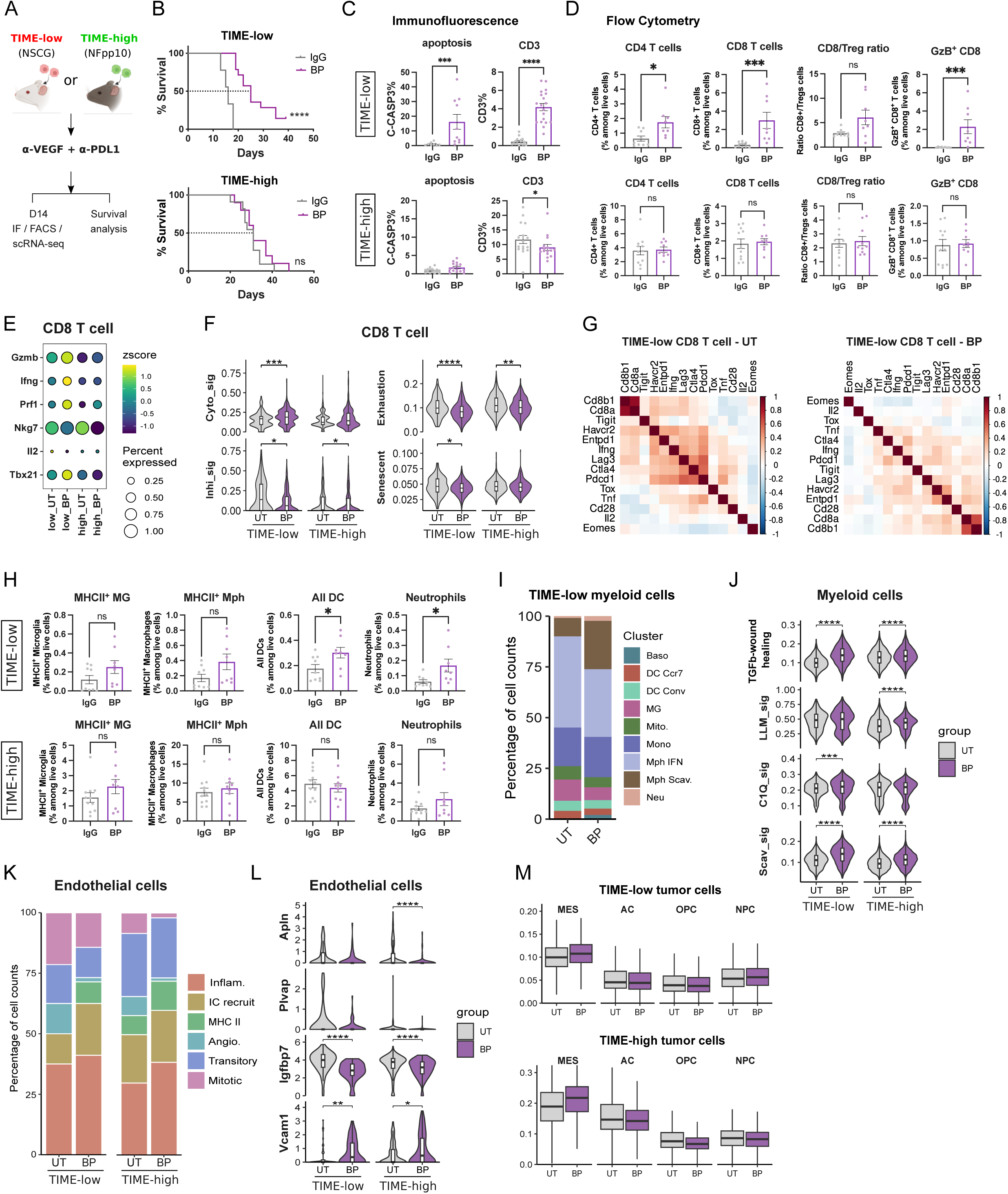
Characterization of T cells and myeloid cells in TIME-low and TIME-high GBMs before and after anti-angiogenic immunotherapies. (A) Experimental scheme for anti-angiogenic immunotherapy (BP: B20 + anti-PD-L1) in TIME-low (NSCG) and TIME-high (NFpp10) GBM models. (B) Kaplan-Meier survival curves for TIME-low and TIME-high tumor-bearing mice treated with IgG control or BP therapy. (C) Quantification of apoptosis and CD3+ T cell infiltration by immunofluorescence staining in TIME-low and TIME-high tumors treated with IgG or BP. (D) Flow cytometry analysis of CD4+ T cells, CD8+ T cells, CD8/Treg ratio, and Gzmb+ CD8+ T cells in TIME-low and TIME-high tumors treated with IgG or BP. (E) Dot plot showing expression of effector molecules in CD8+ T cells from TIME-low and TIME-high tumors before and after BP treatment. (F) Violin plots showing expression of cytotoxicity (Cyto_sig), inhibitory (Inhi_sig), exhaustion, and senescence signatures in CD8+ T cells from TIME-low and TIME-high tumors before and after BP treatment. (G) Correlation analysis of “chronically sensitized” CD8+ T cell signature in TIME-low CD8+ T cells from control and BP-treated tumors. (H) Flow cytometry quantification of MHCII+ microglia, MHCII+ macrophages, dendritic cells (All DC), and neutrophils in TIME-low and TIME-high tumors treated with IgG or BP. (I) Stacked bar plots showing the distribution of myeloid cell subpopulations in TIME-low tumors before (UT) and after BP treatment. (J) Violin plots showing expression of TGFβ-wound healing, lipid-laden macrophage (LLM_sig), C1Q (complement) and Scavenger immunosuppressive signatures in myeloid cells from TIME-low and TIME-high tumors before and after BP treatment. (K) Stacked bar plots showing the distribution of EC subclusters in TIME-low and TIME-high tumors before and after BP treatment. (L) Violin plots showing expression of endothelial cell markers in TIME-low and TIME-high tumors before and after BP treatment. (M) Box plots showing the distribution of tumor cell subtypes in TIME-low and TIME-high tumors before and after BP treatment. Data are presented as mean ± SEM. Statistical significance: *p < 0.05, **p < 0.01, ***p < 0.001, ****p < 0.0001. (Mann-Whitney test or as indicated).

ScRNA-seq of immune cell clusters corroborated IF and FACS findings (**Fig. S3C**). In BP-treated TIME-low tumors, CD8+ T cells showed elevated *Gzmb*, *Ifng*, *Prf1*, and *Tbx21* expression (**Fig. 3E**), and GSEA revealed broad upregulation of T cell proliferation, differentiation, and activation pathways (**Fig. S3D**). TIME-high tumors remained largely unresponsive (**Fig. S3E**). Cytotoxicity signatures^50^ increased while inhibitory^50^, exhaustion^51^, and senescence^52^ signatures decreased in TIME-low T cells; TIME-high T cells showed no significant changes (**Fig. 3F**). Interestingly, CD8+ T cell signature correlations decreased post-treatment in TIME-low tumors, suggesting adaptive resistance (**Fig. 3G**). TIME-high CD8 T cells showed no significant concordance amongst genes of csCD8^+^T-cell signature, and this was also not improved after treatment (**Fig. S3F**).

Myeloid cell responses mirrored T-cell patterns. BP-treated TIME-low tumors showed increased infiltration of MHC II^+^ microglia, macrophages, dendritic cells, and neutrophils, while TIME-high tumors had minimal changes (**Fig. 3H**). Notably, we also observed an increase in immunosuppressive scavenger macrophages and elevated TGF-β/wound healing and complement immunosuppression signatures in TIME-low GBM, potentially explaining relapse (**Fig. 3I-J**). TIME-high tumors already exhibited high immunosuppressive myeloid activity, with modest further increases (**Fig. 3J**). BP therapy improved vascular function in both subtypes, with more immune-recruiting and MHC II^+^ endothelial cells, fewer angiogenic/mitotic ECs (**Fig. 3K**), and gene expression shifts including downregulated *Apln*, *Plvap*, *Igfbp7,* and upregulated *Vcam1* (**Fig. 3L**). However, these vascular changes did not yield survival benefits in TIME-high tumors, paralleling modest bevacizumab effects in patients.

It is noteworthy that the BP treatment caused an increase in mesenchymal-like signatures in both TIME-low and TIME-high tumors (**Fig. 3M**). This shift aligns with previous studies^53–56^ and may reflect a treatment-induced change that contributes to eventual adaptive resistance.

In summary, TIME-low tumors, exhibiting a rather angiogenic and leaky vasculature with exhausted tumor-reactive CD8 T cells, responded transiently to antiangiogenic ICB therapy, reflecting patterns in recurrent human GBM where PD-1 blockade modestly activates T cells but increases myeloid immunosuppression^53^. TIME-high tumors with a low angiogenic index, abundant immunosuppressive myeloid cells and dysfunctional T cells were non-responsive. These findings align with clinical observations that BP is ineffective in human GBM patients^14^, underscoring the need for subtype-specific therapeutic strategies.

### CD40 agonist worsens the survival outcome of TIME-high GBM

Next, we tested a CD40 agonist based on its ability to reprogram macrophages to an immune-stimulating, antigen-presenting state, which has been an effective strategy in several tumor types^57–62^. Notably, CD40 was not only expressed in antigen-presenting cells (APCs) but also in some endothelial cells, while CD40 ligand (*Cd40lg*) was exclusive to T cells (**Fig. S4A**).

CD40Ag had no significant survival benefit in TIME-low GBM-bearing mice, although it modestly increased expression of *Cd80*, *Cd86*, and MHC II genes in both microglia and macrophages (**Fig. 4A-B, 4E**, **S4E-F**). In contrast, CD40Ag alone or combined with DC vaccination, curtailed the survival in TIME-high tumor-bearing mice compared to control (**Fig. 4B**, **S4C**), concomitant with increased microglia infiltration (**Fig. 4C-D**, **S4D**), decreased tumor cell apoptosis (**Fig. S4B**), reduced expression of the APC gene machinery, and heightened *Spp1* (osteopontin) expression in both microglia and macrophages (**Fig. 4E-G**). Interestingly, changes in *Il12*, *Cd80*, *Cd86*, and MHC II expression in dendritic cells were minimal, indicating that the negative treatment effects were predominantly targeting microglia and macrophages (**Fig. S4G**).

**Figure 4.**
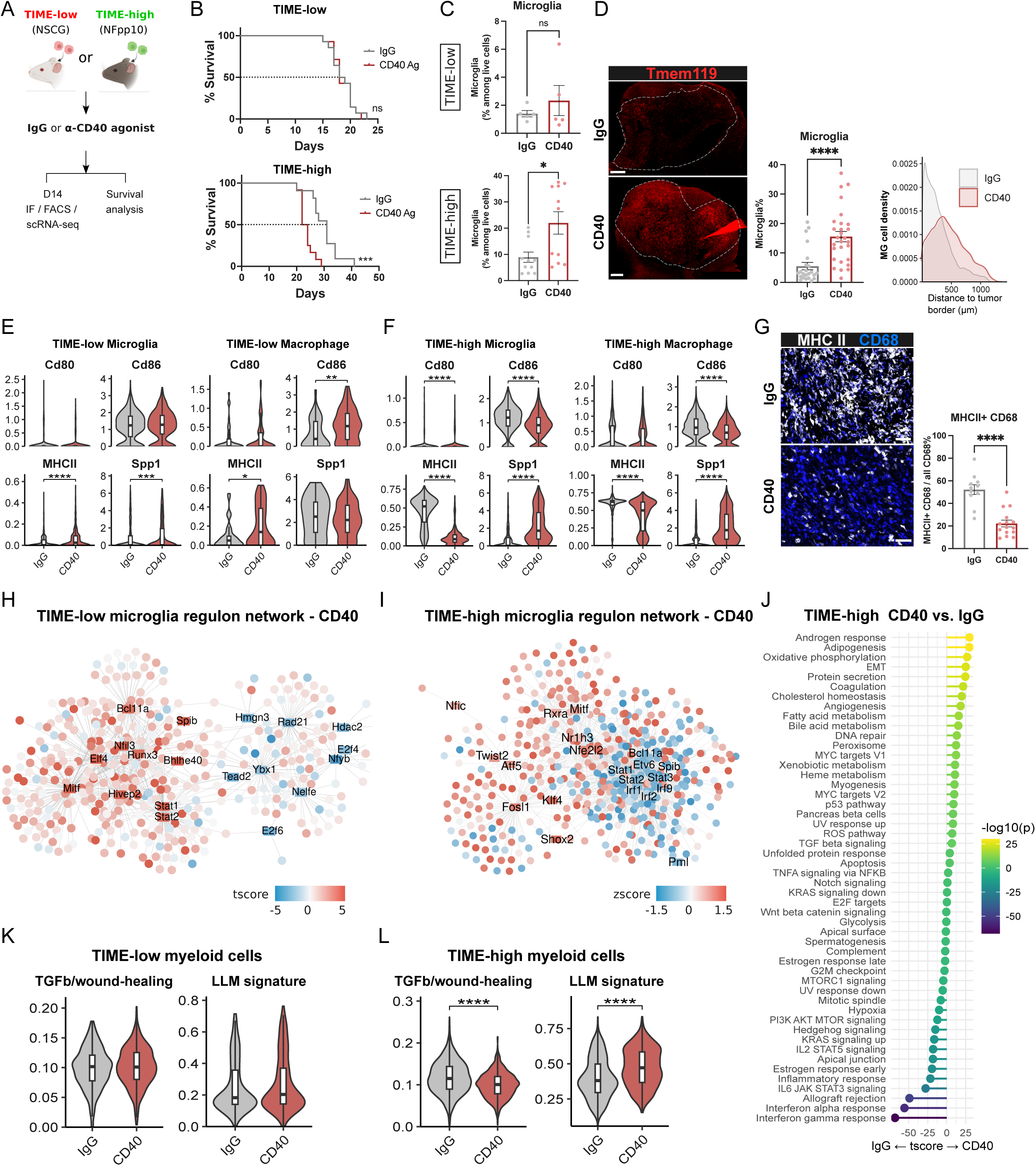
Anti-CD40 agonist treatment worsens the outcome of TIME-high GBMs. (A) Experimental scheme for anti-CD40 agonist treatment in TIME-low and TIME-high GBM models. (B) Kaplan-Meier survival curves for TIME-low and TIME-high tumor-bearing mice treated with IgG control or anti-CD40 agonist. (C) Flow cytometry quantification of microglia in TIME-low and TIME-high tumors before and after anti-CD40 agonist treatment. (D) Representative IF images (left), quantification (middle) and distribution (right) of microglia in TIME-high tumors treated with IgG or anti-CD40 agonist. (E-F) Violin plots showing expression of Cd80, Cd86, and MHC II genes in microglia and macrophages from TIME-low (E) and TIME-high (F) tumors after IgG or anti-CD40 agonist treatment. (G) Representative immunofluorescence images (left) and quantification (right) of MHCII+ CD68+ myeloid cells in TIME-high tumors treated with IgG or anti-CD40 agonist. (H-I) Gene regulatory networks inferred by SCENIC for microglia in TIME-low (H) and TIME-high (I) tumors after anti-CD40 agonist treatment. (J) GSEA of hallmark pathways in myeloid cells from TIME-high tumors comparing anti-CD40 vs. IgG treatment. (K-L) Violin plots showing expression of TGFβ-wound healing and LLM signatures in myeloid cells from TIME-low (K) and TIME-high (L) tumors after IgG or anti-CD40 agonist treatment. Data are presented as mean ± SEM. Statistical significance: *p < 0.05, **p < 0.01, ***p < 0.001, ****p < 0.0001. (Mann-Whitney test or as indicated).

SCENIC analysis^63^ revealed a moderate increase of inflammation- and interferon-related TF activities in TIME-low GBM, whereas CD40Ag in TIME-high tumors downregulated them and rather activated Fosl1, Nfic, and Atf5, which have tumor-promoting activities^64,65^ (**Fig. 4H-J**, **S4H-J**). The suppressive LLM expression signature was significantly increased in myeloid cells of TIME-high tumors, concomitant with the activation of cholesterol and fatty acid metabolism pathways, but was unchanged in TIME-low tumors (**Fig. 4J-L**).

### CD40 agonist induces T cell dysfunction, NK cell reduction, abnormal angiogenesis, and tumor immune evasion in TIME-high GBMs

FACS of the adaptive immune cell populations indicated a marked reduction in NK cells but an increase in CD8+ T cells in TIME-high GBM after CD40 treatment, while CD4^+^ T cells remained relatively unchanged (**Fig. 5A**). CD8+ T cells, however, upregulated *Pdcd1*, *Ctla4*, and *Tox*, and *Gzmk* (**Fig. 5B**), which was also reflected in the largely decreased concordance of CD8^+^ T-cells, reminiscent of a severely dysfunctional state (**Fig. 5C**). Similar but less pronounced trends were observed in TIME-low tumors (**Fig. S5A-C**). Interestingly, NK cells were reduced without altering their apoptotic or proliferative rate (**Fig. 5A, 5D, S5C-E**). CellChat analysis^66^ depicted an interaction of NK cells with endothelial cells via L-selectin (*Sell)-Podxl* (**Fig. S5G-H**), and recruitment signals with TA-Microglia through the *Ccl2/12-Ccr2*, and *Cxcl12-Cxcr4* axes (**Fig. 5E**, **S5I**). CD40Ag significantly reduced the expression of *Ccl2* in microglia and *Cxcl12* in endothelial cells, along with the NK cell receptors *Ccr2* and *Cxcr4* (**Fig. 5F**). Thus, CD40 agonist impaired NK cell recruitment in TIME-high tumors. This reduction likely contributed to the immunosuppressive phenotype because NK cells retained high expression of *Ifng*, *Gzma*, *Gzmb*, *Prf1* independent of treatment (**Fig. S5C**), and depleting NK cells led to a reduced survival trend of TIME-high GBM-bearing mice (**Fig. S5F**).

**Figure 5.**
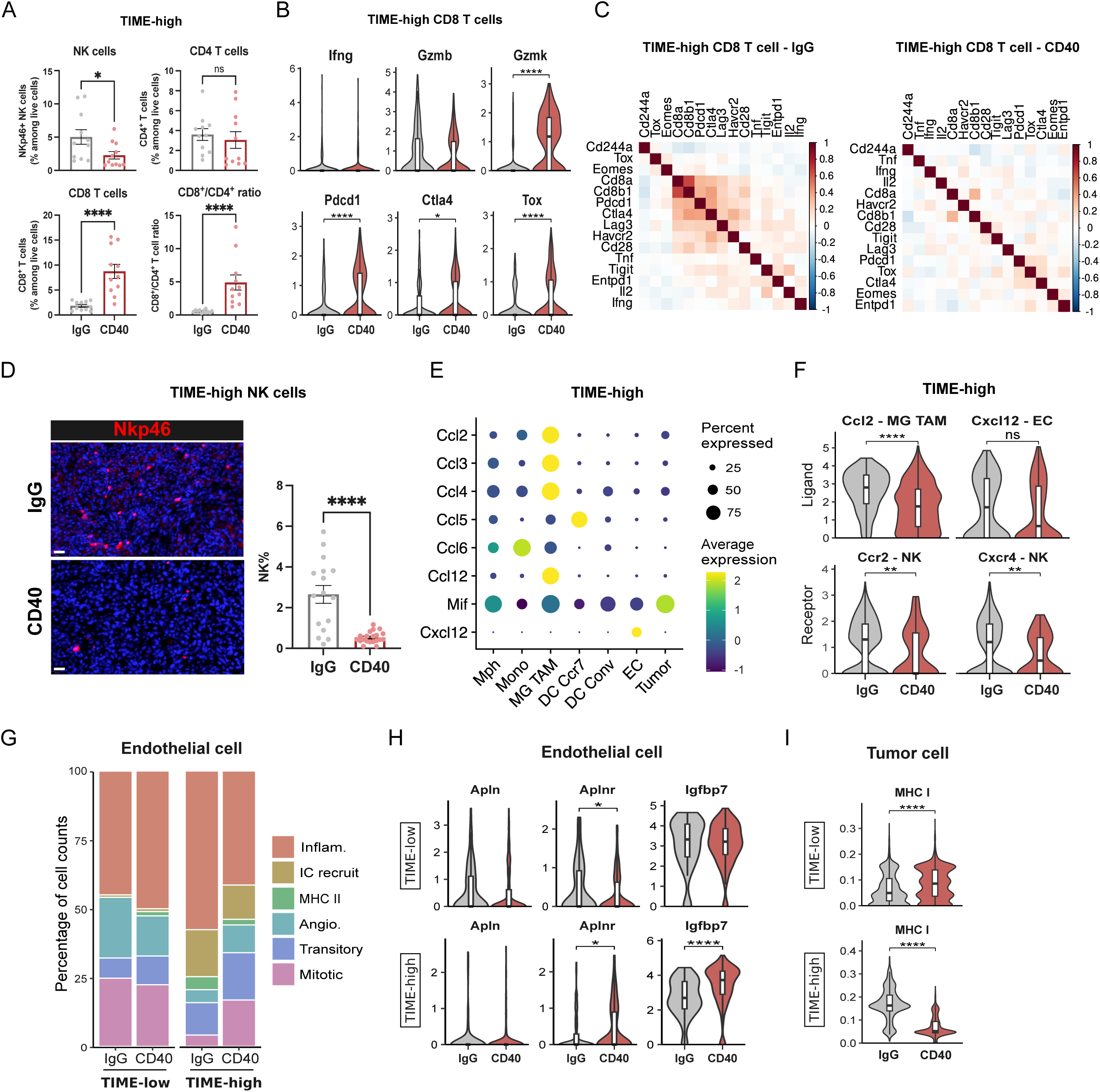
Understanding the resistant mechanisms of TIME-high GBMs to anti-CD40 agonist. (A) Flow cytometry quantification of NK cells, CD8+ T cells, CD4+ T cells, and CD8+/CD4+ T cell ratio in TIME-high tumors treated with IgG or anti-CD40 agonist. (B) Violin plots showing expression of effector molecules and inhibitory receptors in CD8+ T cells from TIME-high tumors treated with IgG or anti-CD40 agonist. (C) Correlation analysis of “chronically sensitized” CD8+ T cell signature in TIME-high tumors treated with IgG or anti-CD40 agonist. (D) Representative immunofluorescence images (left) and quantification (right) of NK cells in TIME-high tumors treated with IgG or anti-CD40 agonist. (E)Dot plot showing expression of chemokines involved in NK cell recruitment across different myeloid cell subpopulations. Color intensity indicates expression level, and dot size represents percentage of cells expressing the gene. (F) Violin plots showing expression of chemokine ligands and receptors in different cell types involved in NK cell recruitment after IgG or anti-CD40 agonist treatment. (G) Stacked bar plots showing the distribution of endothelial cell subclusters in TIME-low and TIME-high tumors treated with IgG or anti-CD40 agonist. (H) Violin plots showing expression of endothelial cell markers in TIME-low and TIME-high tumors treated with IgG or anti-CD40 agonist. (I) Violin plots showing MHC I expression on tumor cells after anti-CD40 agonist treatment in TIME-low and TIME-high tumors. Data are presented as mean ± SEM. Statistical significance: *p < 0.05, **p < 0.01, ***p < 0.001, ****p < 0.0001 (Mann-Whitney test or as indicated).

As ECs also expressed CD40 (**Fig. S4A**), CD40 agonist elevated the proportions of angiogenic and mitotic ECs and decreased inflammatory ECs in TIME-high tumors, but not in TIME-low tumors (**Fig. 5G**). These findings aligned with the upregulation of the angiogenic molecules *Aplnr*, *Igfbp7* in ECs and *Spp1* in TAMs of TIME-high tumors (**Fig. 5H, 4F**). In tumor cells, MHC I expression was significantly decreased in TIME-high tumors following CD40 treatment, while it was moderately increased in TIME-low tumors (**Fig. 5I**). Furthermore, TIME-high tumor cells shifted towards a more AC and NPC-like phenotype, while there were no significant changes in TIME-low tumors post-treatment (**Fig. S5J**).

Overall, CD40 agonist treatment elicited divergent responses between TIME-high and TIME-low GBMs. TIME-high GBM became more immunosuppressive and angiogenic with an abbreviated survival while TIME-low GBM showed trends of an immune response without a survival benefit.

### IPI-145 enhances immune responses to anti-angiogenic immunotherapies in TIME-high GBMs

We next investigated the PI3Kγ/δ inhibitor IPI-145, given that PI3Kγ activity promotes immunosuppression in TAMs by inhibiting NFκB and activating C/EBPβ^67^, and was shown to be upregulated as an adaptive resistance mechanism to antiangiogenic therapy^68^. Congruently, BP treatment increased *Pik3cg* and *Pik3cd* expression, PI3K-Akt-mTOR signaling, and C/EBPβ transcriptional activity in myeloid cells in both TIME-low and TIME-high tumors (**Fig. S6A-C**). This was accompanied by elevated *Arg1* and *Il1b* (**Fig. S6C**), indicative of a shift toward an immunosuppressive myeloid phenotype. Importantly, the combination of IPI-145 and BP (BPI) substantially extended the median survival in TIME-high tumor-bearing mice, while TIME-low tumors showed no additional survival benefit from IPI-145 (**Fig. 6A, 6B**). FACS revealed that BPI did not affect CD45^+^ cells and MHCII^+^ macrophage levels in TIME-low tumors, but increased MHCII^+^ microglia and reduced neutrophils (**Fig. 6C**, **S6D**). TIME-high tumors also displayed elevated CD45^+^ and MHCII^+^ microglia levels upon BPI with reduced neutrophils, while MHCII^+^ macrophages remained unchanged (**Fig. 6C**, **S6D**). Transcriptional profiling of BPI-treated TIME-low tumors revealed upregulation of *Cd86* and MHCII in both microglia and macrophages, suggesting activation (**Fig. 6D**, **S6E**). Notably, *Spp1* expression decreased in macrophages, but remained unchanged in microglia (**Fig. 6D**, **S6E**). In TIME-high tumors, MHCII+ myeloid cell levels were not altered, accompanied by a marginal increase in OPN expression (**Fig. 6E**).

**Figure 6.**
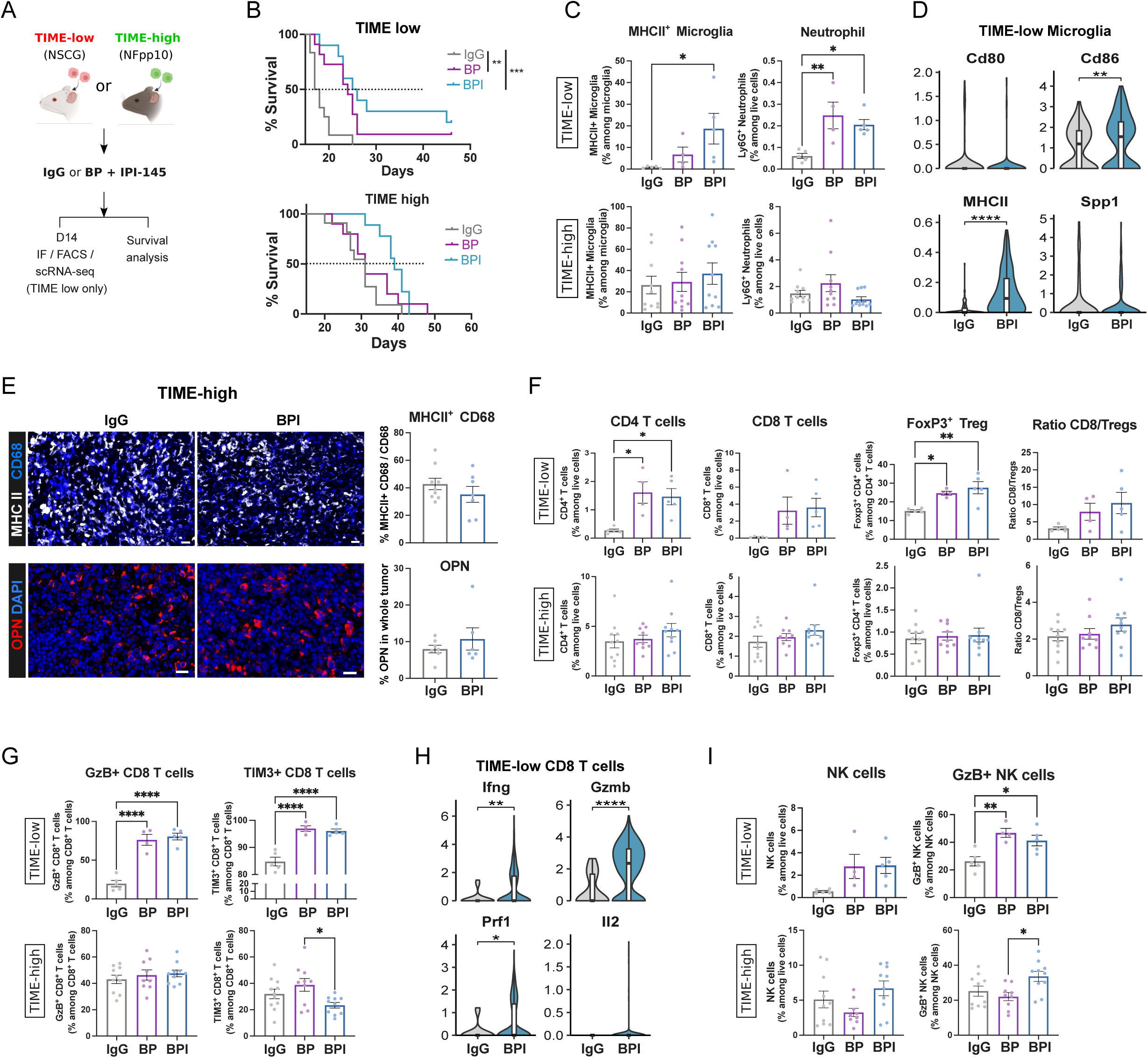
Myeloid repolarization in TIME-high GBMs with IPI-145 improves response to anti-angiogenic immunotherapies. (A) Experimental scheme for BP and PI3K inhibitor (IPI-145) combination treatment (BPI) in TIME-low and TIME-high tumors. (B) Kaplan-Meier survival curves for TIME-low and TIME-high tumor-bearing mice treated with IgG, BP, or BPI. (C) Flow cytometry quantification of MHCII+ microglia and neutrophils in TIME-low and TIME-high tumors treated with IgG, BP, or BPI. (D) Violin plots showing expression myeloid markers in microglia from TIME-low tumors treated with IgG or BPI. (E) Representative immunofluorescence images and quantification of MHCII+ CD68+ myeloid cells and OPN expression in TIME-high tumors treated with IgG or BPI. (F) Flow cytometry quantification of CD4+ T cells, CD8+ T cells, FoxP3+ Tregs and CD8/Treg ratio in TIME-low and TIME-high tumors treated with IgG, BP, or BPI. (G) Flow cytometry quantification of GzB+ CD8+ T cells and TIM3+ CD8+ T cells in TIME-low and TIME-high tumors treated with IgG, BP, or BPI. (H) Violin plots showing expression of effector molecules in CD8+ T cells from TIME-low tumors treated with IgG or BPI. (I) Flow cytometry quantification of NK cells and GzB+ NK cells in TIME-low and TIME-high tumors treated with IgG, BP, or BPI. Data are presented as mean ± SEM. Statistical significance: *p < 0.05, **p < 0.01, ***p < 0.001, ****p < 0.0001. (Mann-Whitney test or as indicated).

T cell responses also differed by TIME status. In TIME-low tumors, CD4^+^, CD8^+^, and Treg infiltration was similar between BP and BPI, though the CD8/Treg ratio trended upward with BPI (**Fig. 6F**). TIME-high tumors exhibited increased CD4^+^ and CD8^+^ T cell infiltration without a rise in Tregs, resulting in a consistently higher CD8/Treg ratio (**Fig. 6F**). In TIME-low tumors, BPI increased GzB^+^, TIM3^+^, and PD1^+^ CD8^+^ T cells compared to IgG, but not BP (**Fig. 6G**, **S6F**). scRNA-seq revealed elevated expression of *Ifng, Gzmb,* and *Prf1*, along with checkpoint receptors *Pdcd1, Ctla4,* and *Havcr2*, but reduced *Tox*, diverging from the CD40 agonist response (**Fig. 6H**, **S6G**). In TIME-high tumors, BPI treatment did not raise GzB^+^ CD8^+^ T cells, but reduced TIM3^+^ cells with a downward trend in PD1^+^ cells (**Fig. 6G**, **S6F**), suggesting less exhaustion.

NK cell dynamics mirrored T cell trends. In TIME-low tumors, NK infiltration increased with BP and BPI, though GzB^+^ NK cells were slightly lower with BPI. In TIME-high tumors, BPI treatment significantly increased the infiltration of GzB^+^ NK cells (**Fig. 6I**)-a sharp contrast to the response seen with the CD40 agonist. Overall, BPI enhances cytotoxic CD8^+^ T and NK cell responses in TIME-high tumors, likely driving the improved therapeutic effect in this context.

## Discussion

Despite transformative advances in immunotherapy, GBMs continue to defy most interventions, with only a small fraction of patients experiencing beneficial responses. This persistent resistance likely reflects not just the genetic and phenotypic heterogeneity within tumor cells themselves, but a broader variability across the TIME—one that remains difficult to predict and even harder to modulate. To address this, we leveraged scRNA-seq and spatial proteomics to construct a more granular view of GBM TIME composition. From these data emerged three reproducible TIME archetypes—TIME-low, TIME-med, and TIME-high— each distinguished by distinct patterns of immune infiltration, and vascular and functional immune states. Crucially, these were not merely descriptive categories. By identifying and validating murine models that faithfully mirrored each subtype of human GBM, we were able to observe striking differences in immunotherapy responses and begin to parse the mechanisms that underlie therapeutic resistance.

Our observations confirmed and expanded upon trends previously reported in bulk RNA-seq studies^15^, but may also seem cautionary in how we interpret prior bulk data. TIME-low tumors, for instance, exhibit the expected “immune desert” phenotype: sparse presence of T cells and few macrophages, and a leaky vasculature. TIME-high tumors, by contrast, are heavily immune-infiltrated—yet this apparent abundance conceals a deeply suppressive environment dominated by lipid-laden, scavenger-type macrophages and severely dysfunctional T cells. TIME-med tumors occupy a more ambiguous space: their vasculature is highly angiogenic, but their immune compartment is relatively active, marked by both activated T cells and immune-stimulatory myeloid cells.

These distinctions carry clinical relevance. TIME-low tumors, while initially responsive, tended to relapse, undergoing a phenotypic drift toward a TIME-high state. TIME-high tumors, meanwhile, remained largely unresponsive, challenging the assumption that more immune infiltration is necessarily beneficial. In contrast, TIME-med tumors showed more favorable responses to ICBs and anti-angiogenic agents. Notably, patients with TIME-med-like tumors also exhibited improved outcomes following ICB combined with oncolytic virotherapy^8^— suggesting a degree of immunologic pliability that TIME-low and TIME-high tumors seem to lack. This prompts the question, if approximately 30% of patients develop TIME-med GBM, why do not more respond to immunotherapies? One possibility is that standard chemotherapy and radiotherapy, as demonstrated in preclinical models, may alter the TIME status, thereby reducing the proportion of patients with TIME-med GBM. This raises the intriguing possibility that, for such patients, ICB or other immunotherapies might be administered before conventional treatments—a strategy shown to be beneficial in several different cancer types^69,70^. Supporting this concept, a recent case report of neoadjuvant triplet ICB in newly diagnosed GBM showed promising results, with no evidence of recurrence at 17 months, prompting the launch of a clinical trial in November 2025^71^.

Our findings further underscore the necessity of carefully selecting appropriate preclinical models to mirror human GBM TIME subtypes. While the widely used GL261 model reflects some of the phenotypes of human TIME-med tumors and displays a more robust immunotherapy responsiveness^72–74^, it is notably a carcinogen-induced GBM and does not capture the complexities of TIME-low or TIME-high environments. By incorporating genetically-engineered NSCG and NFpp10 models, we established faithful murine counterparts for TIME-low and TIME-high GBMs, respectively, and thereby were able to reproduce better the spectrum of immune phenotypes seen in patients. At the single-cell level, these models revealed clear divergence in both quantitative and functional myeloid and lymphocyte characteristics. A significant insight from our study concerns the complex, often paradoxical, effects of targeting TAMs, revealing that efforts to reprogram them therapeutically can backfire. While CD40 agonists have shown promise in other malignancies by driving TAMs toward pro-inflammatory states^62,75–79^, their effects in GBM were markedly context-dependent. In TIME-low tumors, CD40 activation was insufficient to improve outcomes, and actually worsened outcomes in TIME-high tumors by amplifying the suppressive, scavenger-like macrophage phenotype, reducing NK cell infiltration, and deepening immune dysfunction.

These results align with some preclinical reports, which noted increased T cell infiltration but limited functionality after CD40 treatment^60,61^. What our data suggest is that CD40 agonists may have detrimental effects in some GBM subtypes—a consideration that feels increasingly urgent as these agents move through clinical trials^57^. They also stress the critical importance of TIME-based patient stratification and the need for predictive biomarkers to identify those most likely to benefit.

We also tested whether targeting PI3K signaling, a known mediator of immune suppression in myeloid cells, could mitigate the resistance observed in TIME-high tumors. Combining the PI3Kγ/δ inhibitor IPI-145 with anti-angiogenic immunotherapy extended median survival and partially enhanced cytotoxic immune function, likely constrained by ongoing TAM plasticity and persistent vascular dysfunction. Inhibiting myeloid PI3K signaling appeared to nudge the system in the right direction, but was not sufficient to reprogram suppressive macrophages in a durable manner. These data suggest that TIME-high GBMs require distinct immunomodulating strategies that more efficiently reprogram the myeloid compartment to elicit immune responses.

In summary, this work outlines a conceptual and practical framework for tailoring immunotherapy in GBM. TIME-med tumors appear uniquely poised to respond, while TIME-low and TIME-high tumors resist through distinct, yet equally entrenched, forms of TAM-mediated suppression. These findings argue against a one-size-fits-all model of immunotherapy and call instead for subtype-specific strategies, ideally anchored in reliable biomarkers and preclinical models that do not oversimplify the immune landscape. Whether such precision approaches can overcome adaptive resistance remains to be seen, but the alternative, treating all GBMs as immunologically equivalent, seems increasingly untenable.

## Ethics

For the MILAN study, the electronic medical records of glioblastoma patients who were treated at the University Hospitals Leuven (Belgium), Ziekenhuis Oost-Limburg Genk (Belgium) and MUMC+ Maastricht (The Netherlands) between 12/2003 and 07/2021 were reviewed. Data collection and analysis were carried out in accordance with data protection guidelines. The study was approved by the medical ethics review board of all participating hospitals (*S59804, S61081, S62248, 18/0021R, 19/0021R, METC 16-4-022*). Written informed consent is obtained by all patients. Animal procedures were approved by the Institutional Animal Care and Research Advisory Committee of the KU Leuven (ECD 192/ 2019) and were performed following the institutional and national guidelines and regulations.

## Acknowledgments

We kindly thank Kevin Feyen, Martine Nijs, Polina Zahdai, Nena Dupont, Nuray Bögürcü-Seidel for their technical assistance. We gratefully acknowledge the VIB Flow Core, the VIB Single Cell Core, and the VIB Nucleomics Core in Leuven. We are grateful to Prof. Michael Platten for providing the GL261 scRNA-seq data and for the invaluable discussion.

## Author contributions

L.W, M.G and G.B designed most experiments. L.W, M.G, M.D, C.D and C.P-M performed all experiments unless specified. M.V, P.N and F.D.S performed and analyzed MILAN multiplex immunohistochemistry. L.W and K.S.G performed bioinformatic analysis. A.G, K.W, A.B and D.L provided critical input. G.B conceptually planned and supervised the study. L.W wrote the initial draft of the manuscript; L.W, K.S.G and G.B edited the manuscript.

## Data availability

The newly generated murine scRNA-seq data (fastq files and processed data) have been deposited in GEO under accession number GSE301073. The publicly available human scRNA-seq data can be accessed under EGA/GEO accession numbers: EGAS00001003845, EGAS00001004422, EGAS00001005300, EGAS00001004656, EGAS00001004871, GSE182109. The publicly available GL261 murine model scRNA-seq data can be accessed via ArrayExpress: E-MTAB-12523.

---

**Figure S1.**
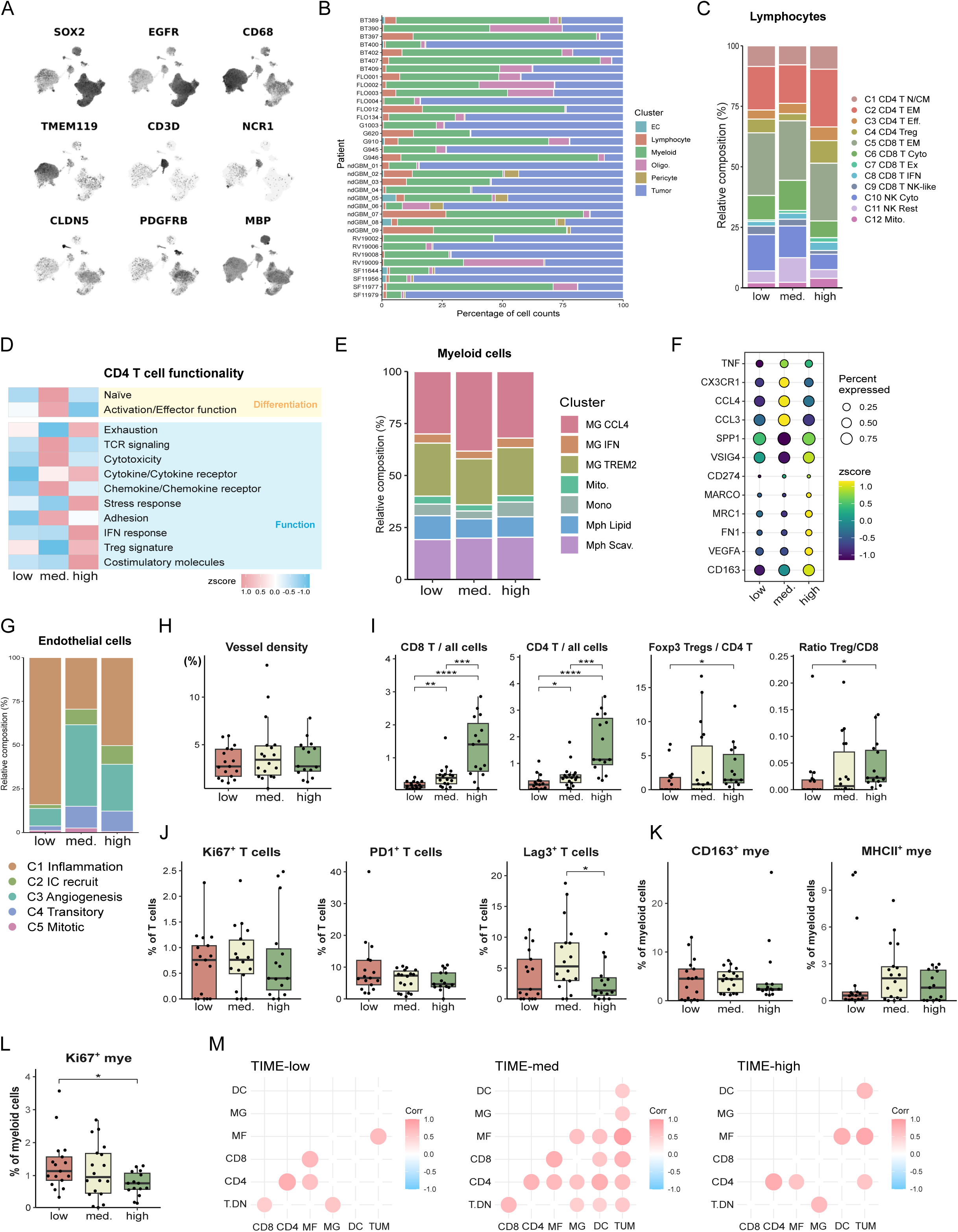
General characterization of human scRNAseq datasets. (A) UMAP plots showing expression of marker genes for major cell types identified in human GBM samples. (B) Stacked bar plot showing the distribution of cell types across different patient samples. (C) Relative composition (%) of lymphocyte subpopulations across TIME-low, TIME-med., and TIME-high samples. (D) Heatmap showing differential expression of CD4+ T cell functional signatures across TIME-low, TIME-med, and TIME-high samples, grouped by differentiation and function categories. (E) Relative composition (%) of myeloid cell subclusters across TIME subtypes. (F) Dot plot showing expression of inflammatory and immuosuppressive markers across TIME subtypes. Color indicates expression level (z-score), and dot size represents percentage of cells expressing the gene. (G) Relative composition (%) of endothelial cell subclusters across TIME subtypes. (H) Vessel density quantification across TIME subtypes. (I) Quantification of CD8 T cells, CD4 T cells, Foxp3+ Tregs as a proportion of CD4 T cells, and Treg/CD8 ratio across TIME subtypes. (J) Percentages of Ki67+, PD1+, and Lag3+ T cells across TIME subtypes. (K) Percentages of CD163+ and MHCII+ myeloid cells across TIME subtypes. (L) Percentages of Ki67+ myeloid cells across TIME subtypes. (M) Correlograms showing cellular interactions between different immune cell populations (T.DN, CD4, CD8, MF, MG, DC) and tumor cells (TUM) in TIME-low, TIME-med, and TIME-high tumors. Statistical significance: *p < 0.05, **p < 0.01, ***p < 0.001, ****p < 0.0001.

**Figure S2.**
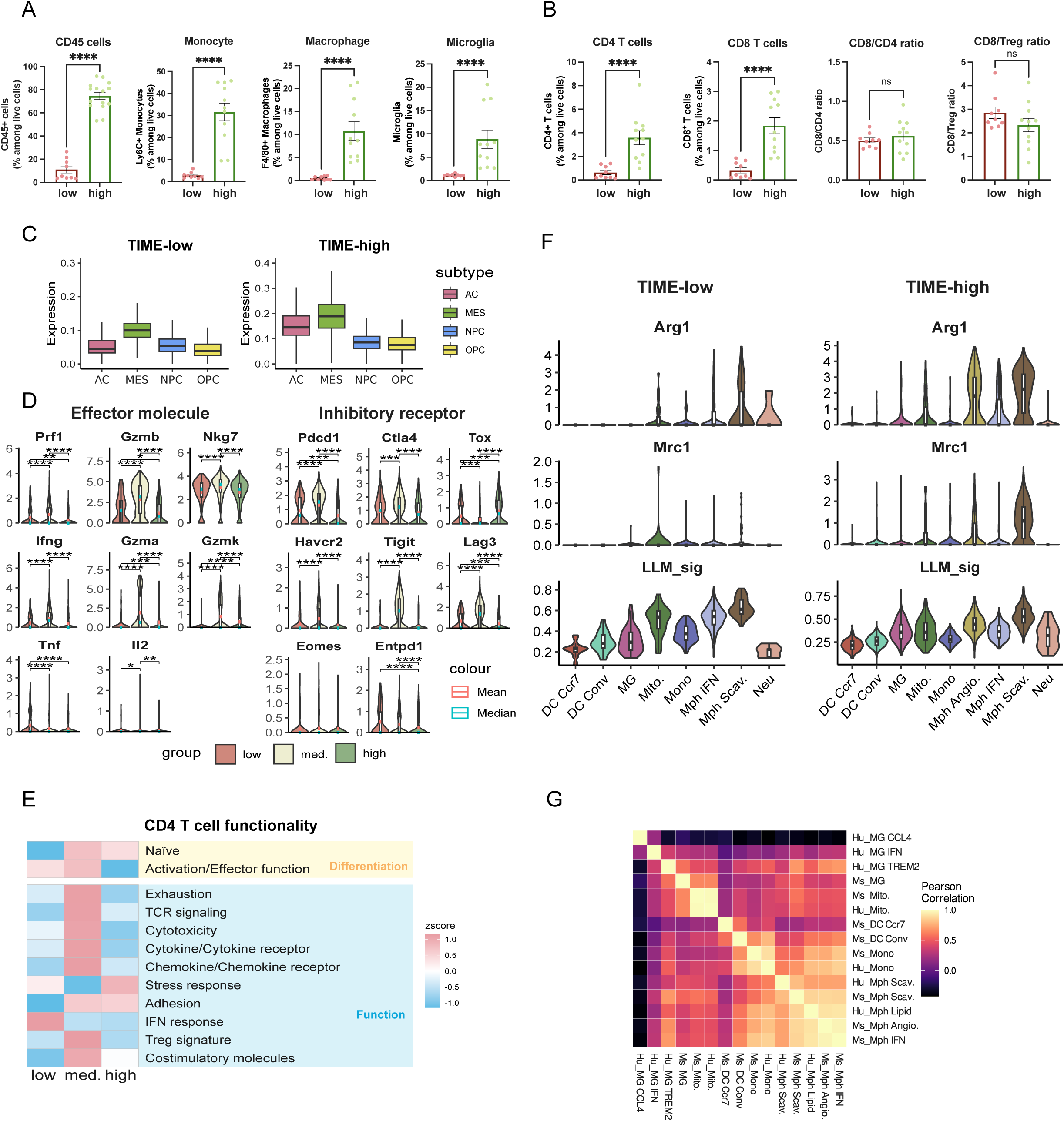
Detailed characterization of immune cell populations in murine GBM models. (A) Flow cytometry quantification of CD45+ immune cells, Ly6C+ monocytes, F4/80+ macrophages, and microglia in TIME-low and TIME-high tumors. (B) Flow cytometry analysis of CD4+ T cells, CD8+ T cells, CD8/CD4 ratio, and CD8/Treg ratio in TIME-low and TIME-high tumors. (C) Distribution of tumor subtypes across TIME-low and TIME-high tumors. (D) Violin plots showing expression of effector molecules and inhibitory receptors in CD8+ T cells from TIME-low and TIME-high tumors. (E) Heatmap showing differential expression of CD4+ T cell functional signatures across TIME-low, TIME-med, and TIME-high murine tumors, grouped by differentiation and function categories. (F) Violin plots showing expression of Arg1, Mrc1, and lipid-laden macrophage (LLM) signature across myeloid subpopulations in TIME-low and TIME-high tumors. (G) Correlation heatmap showing relationships between myeloid cell subpopulations in human and murine tumors. Data are presented as mean ± SEM. Statistical significance: *p < 0.05, **p < 0.01, ***p < 0.001, ****p < 0.0001 (Mann-Whitney test).

**Figure S3.**
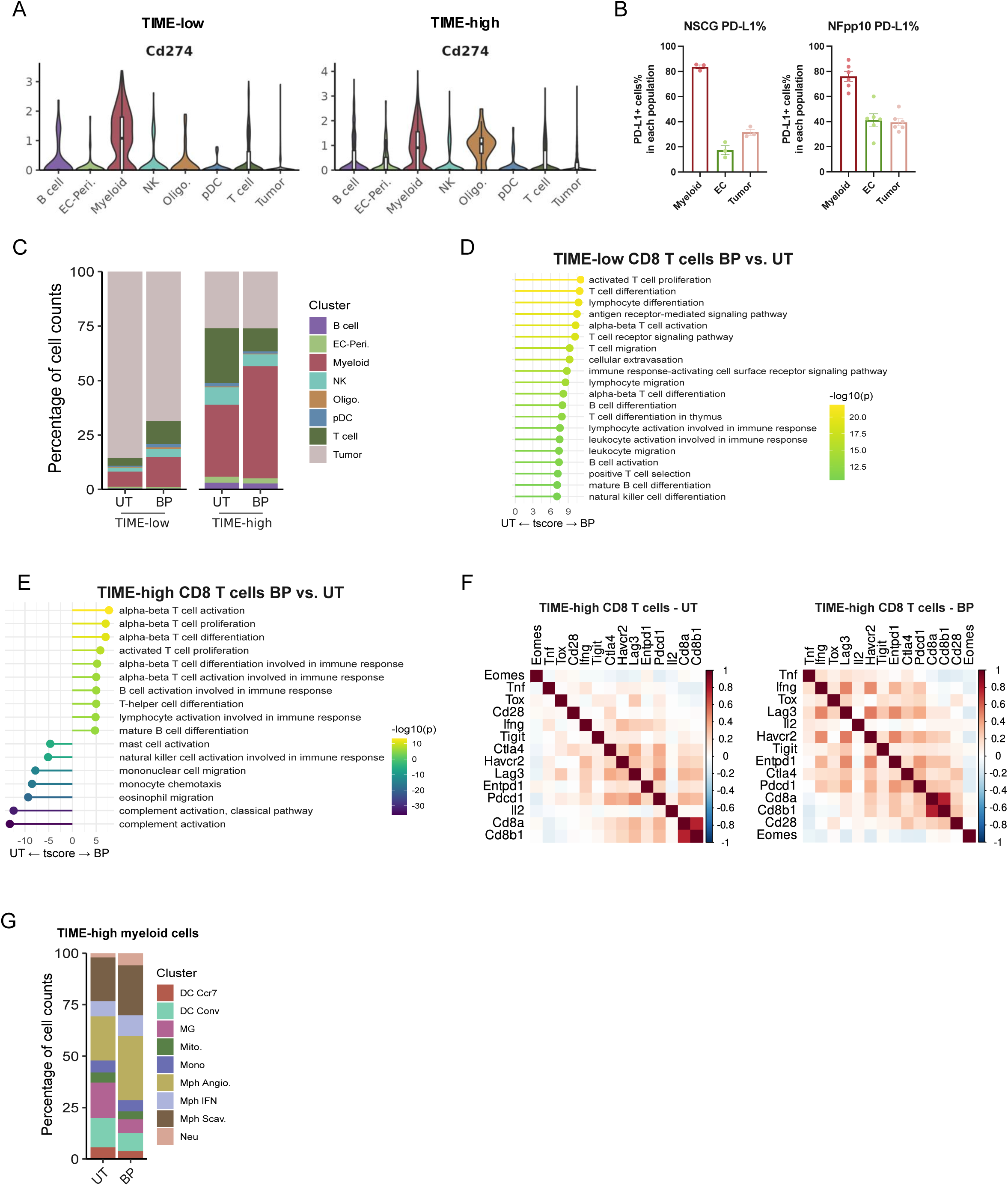
Characterization of T cells and myeloid cells in TIME-low and TIME-high GBMs before and after anti-angiogenic immunotherapies. (A) Violin plots showing expression of Cd274 (PD-L1) across different cell types in TIME-low and TIME-high tumors. (B) Quantification of PD-L1 expression proportion in different cell types of TIME-low and TIME-high tumors. (C) Stacked bar plots showing the proportion of major cell types in TIME-low and TIME-high tumors before (UT) and after BP treatment, based on scRNA-seq data. (D-E) Gene set enrichment analysis (GSEA) of CD8+ T cells in TIME-low and TIME-high tumors comparing BP vs. UT groups using immune-related GO pathways. (F) Correlation analysis of “chronically sensitized” CD8+ T cell signature in TIME-high CD8+ T cells from control and BP-treated tumors. (G) Stacked bar plots showing the distribution of myeloid cell subpopulations in TIME-high tumors before (UT) and after BP treatment. Statistical significance: *p < 0.05, **p < 0.01, ***p < 0.001, ****p < 0.0001.

**Figure S4.**
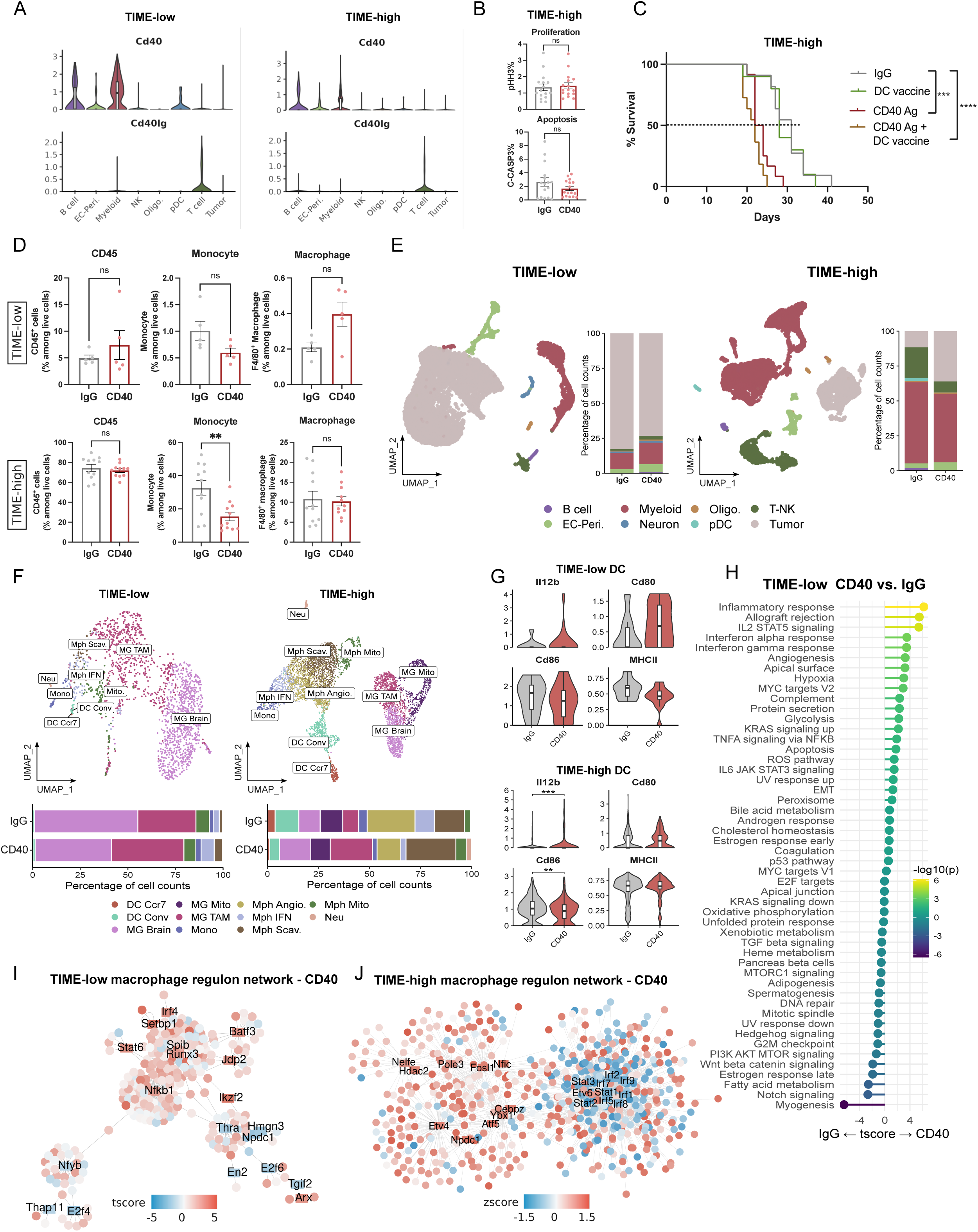
Detailed characterization of anti-CD40 agonist treatment effects in TIME-low and TIME-high GBM models. (A) Violin plots showing expression of CD40 and CD40 ligand (CD40lg) across different cell types in TIME-low and TIME-high tumors. (B) Quantification of tumor cell proliferation and apoptosis in TIME-high tumors treated with IgG or anti-CD40 agonist. (C) Kaplan-Meier survival curves for TIME-high tumor-bearing mice treated with IgG, DC vaccine, anti-CD40 agonist, or anti-CD40 agonist + DC vaccine combination therapy. (D) Flow cytometry quantification of CD45+ cells, monocytes, and macrophages in TIME-low and TIME-high tumors treated with IgG or anti-CD40 agonist. (E) UMAP plots (left) and stacked bar plots (right) showing major cell types identified by scRNA-seq in TIME-low and TIME-high tumors treated with IgG or anti-CD40 agonist. (F) UMAP plots (top) and stacked bar plots (bottom) of myeloid cell subpopulations in TIME-low and TIME-high tumors treated with IgG or anti-CD40 agonist. (G) Violin plots showing expression of Il12, Cd80, Cd86, and MHC II genes in dendritic cells from TIME-low and TIME-high tumors treated with IgG or anti-CD40 agonist. (H) GSEA of hallmark pathways in TIME-low myeloid cells comparing anti-CD40 vs. IgG treatment. (I-J) Gene regulatory networks inferred by SCENIC for macrophages in TIME-low (I) and TIME-high (J) tumors after anti-CD40 agonist treatment. Data are presented as mean ± SEM. Statistical significance: *p < 0.05, **p < 0.01, ***p < 0.001, ****p < 0.0001 (Mann-Whitney test or as indicated).

**Figure S5.**
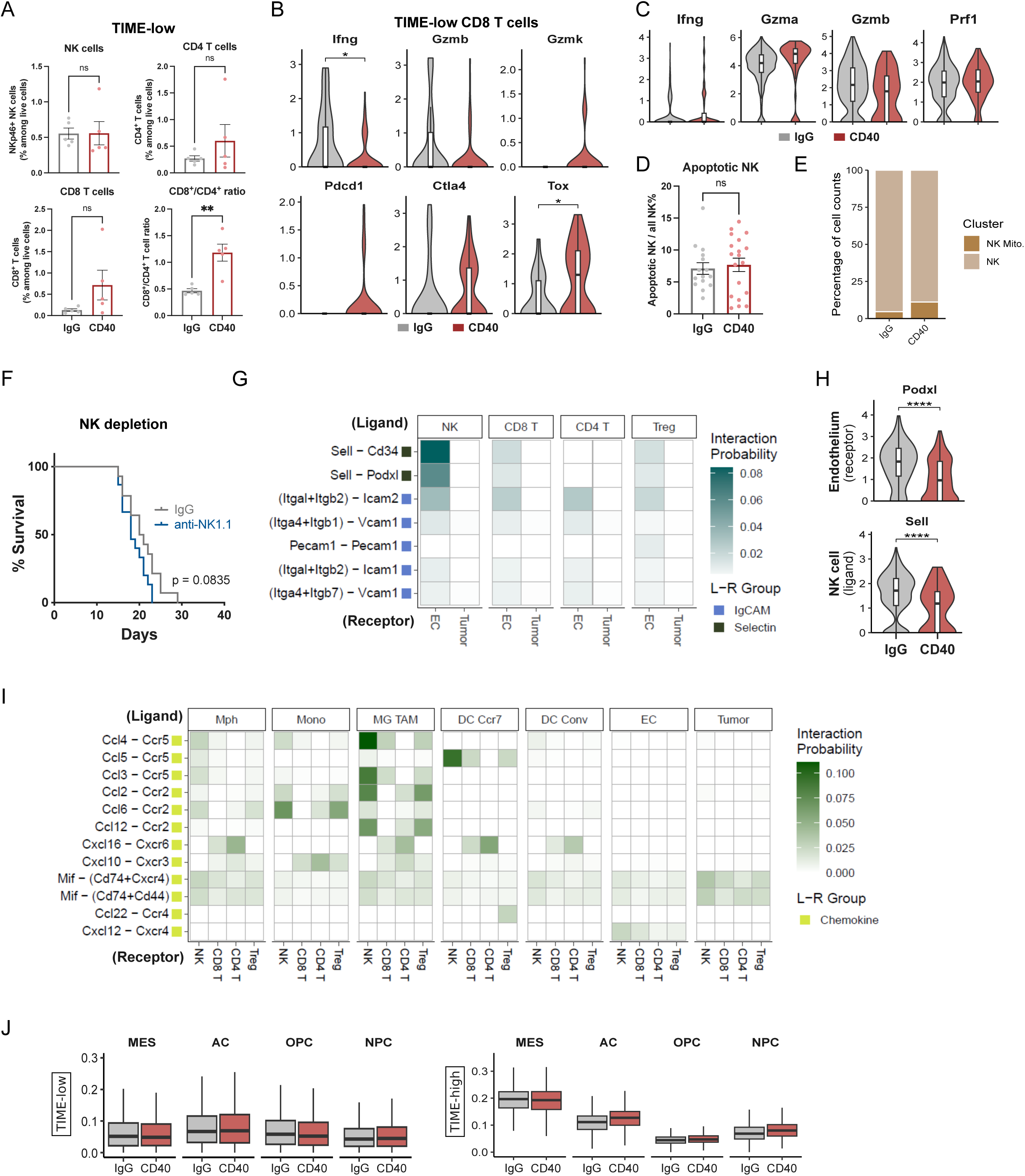
Understanding the resistant mechanisms of TIME-high GBMs to anti-CD40 agonist. (A) Flow cytometry quantification of NK cells, CD4+ T cells, CD8+ T cells, and CD8+/CD4+ T cell ratio in TIME-low tumors treated with IgG or anti-CD40 agonist. (B) Violin plots showing expression of effector molecules and inhibitory receptors in CD8+ T cells from TIME-low tumors treated with IgG or anti-CD40 agonist. (C) Violin plots showing expression of cytotoxic markers in NK cells from TIME-high tumors treated with IgG or anti-CD40 agonist. (D) Quantification of apoptotic NK cells in TIME-high tumors treated with IgG or anti-CD40 agonist. (E) Stacked bar plot showing the proportion of NK cell subpopulations (NK, NK Mito.) in TIME-high tumors treated with IgG or anti-CD40 agonist. (F) Kaplan-Meier survival curves for TIME-high tumor-bearing mice treated with IgG control or anti-NK1.1 therapy. (G) Heatmaps showing expression of ligand (lymphoid cells) and receptor (ECs) interactions involved in NK cell recruitment. (H) Violin plots showing expression of receptor on endothelial cells and ligand on NK cells after IgG or anti-CD40 agonist treatment involved in NK cell recruitment. (I) Heatmaps showing expression of ligand (myeloid cells) and receptor (lymphoid cells) interactions involved in NK cell recruitment. (J) Box plots showing the distribution of tumor cell subtypes in TIME-low and TIME-high tumors treated with IgG or anti-CD40 agonist. Data are presented as mean ± SEM. Statistical significance: *p < 0.05, **p < 0.01, ***p < 0.001, ****p < 0.0001. (Mann-Whitney test or as indicated).

**Figure S6.**
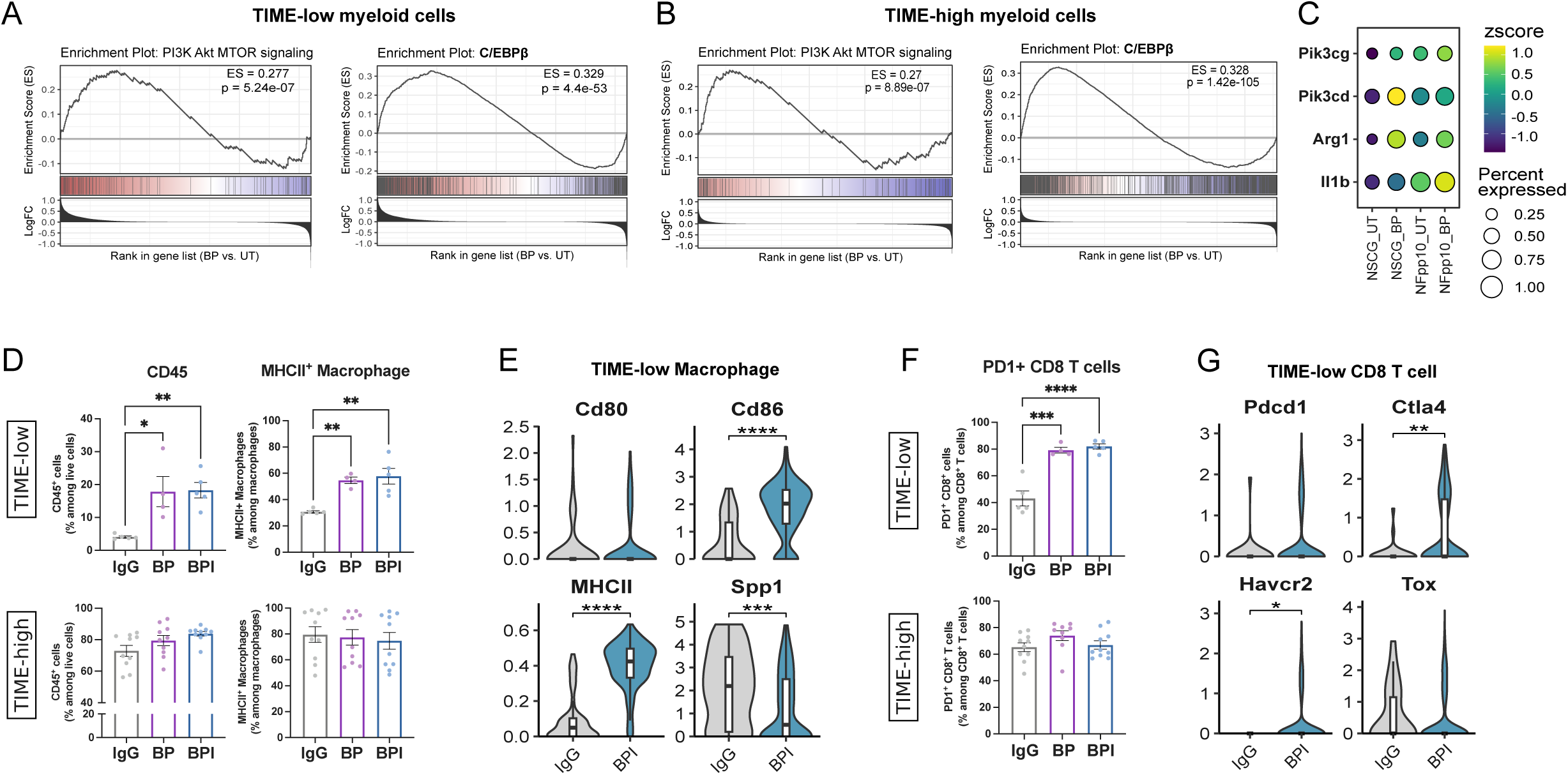
Myeloid repolarization in TIME-high GBMs with IPI-145 improves response to anti-angiogenic immunotherapies. (A-B) GSEA plots showing changes in transcription factor activity for PI3K-Akt-mTOR pathway and C/EBPβ in myeloid cells from TIME-low and TIME-high tumors comparing BP vs. UT groups. (C) Dot plot showing expression of Pik3cg, Pik3cd, Arg1, and Il1b in myeloid cells across different treatment conditions. Color intensity indicates expression level (z-score), and dot size represents percentage of cells expressing the gene. (D) Flow cytometry quantification of CD45+ cells and MHCII+ macrophages in TIME-low and TIME-high tumors treated with IgG, BP, or BPI. (E) Violin plots showing expression of myeloid markers in macrophages from TIME-low tumors treated with IgG or BPI. (F) Flow cytometry quantification of PD1+ CD8+ T cells in TIME-low and TIME-high tumors treated with IgG, BP, or BPI. (G) Violin plots showing expression of inhibitory receptors (Pdcd1, Ctla4, Havcr2, Tox) in CD8+ T cells from TIME-low tumors treated with IgG or BPI. Data are presented as mean ± SEM. Statistical significance: *p < 0.05, **p < 0.01, ***p < 0.001, ****p < 0.0001. (Mann-Whitney test or as indicated).

